# Threshold concentration and random collision determine the growth of the phase-separated huntingtin inclusion from a stable core

**DOI:** 10.1101/2020.08.05.237073

**Authors:** Sen Pei, Theresa C. Swayne, Jeffrey F. Morris, Lesley Emtage

**Affiliations:** Biology Department, City University of New York, York College, Queens, NY; Herbert Irving Comprehensive Cancer Center, Columbia University, New York, NY; Department of Chemical Engineering, The City College of New York, New York, NY; Benjamin Levich Institute, New York, NY; Molecular, Cellular and Developmental Biology Program, City University of New York Graduate Center, New York, NY

## Abstract

The processes underlying formation and growth of unfolded protein inclusions are relevant to neurodegenerative diseases. In *S. cerevisiae*, inclusion bodies formed by mutant huntingtin have characteristics of phase-separated compartments: they are mobile, ovoid, and the contents are diffusible. We have used molecular genetics and quantitative confocal microscopy to probe the relationship between concentration and inclusion growth in vivo. Our analysis and modeling of the growth of mutant huntingtin inclusion bodies (mHtt IBs) suggests that there is a cytoplasmic threshold concentration that triggers the formation of an IB, regardless of proteasome capacity, and that reduction in cytoplasmic mHtt causes IBs to shrink. These findings confirm that the IB is a phase-separated compartment that continuously exchanges material with the cytoplasm. The growth rate of the IB is most consistent with a model in which material is incorporated through collision with the IB. A small remnant of the IB is relatively long-lasting, suggesting that the IB contains a core that is structurally distinct, and which may serve to nucleate it.

## Introduction

Deposits of unfolded protein are characteristic of neurodegenerative disorders, including Huntington’s disease (HD). Visible deposits in the brains of patients with HD are predominantly composed of an N-terminal cleavage product of mutant huntingtin (mHtt) protein (DiFiglia et al., 1997; Lunkes et al., 2002). Causative mutations expand a polyglutamine (polyQ) repeat tract near the N-terminus; in mice, expression of N-terminal fragments of mHtt leads to accumulation in cytoplasmic and nuclear inclusions, recapitulating many aspects of HD (Carty et al., 2015; Hackam et al., 1999; Slow et al., 2003).

While mHtt is expressed in most tissues throughout life, cell death due to mutant Htt is restricted to particular regions of the brain. Most cells are able to cope with persistent expression of moderate, or even high levels of mHtt protein, despite its intrinsic instability. In order to study the response of cells to a constant burden of unstable protein, we have expressed Htt fused GFP from the constitutive GAPDH promoter in *S. cerevisiae*.

When expressed in cultured cells, exon 1 of mHtt, fused to a fluorescent protein or visualized with an antibody, forms cytoplasmic inclusions; often there is initially only one, ovoid inclusion per cell (Arrasate et al., 2004; Yamamoto et al., 2006). In yeast, mutant Htt(exon1)-GFP also typically forms a singular, ovoid inclusion, along with small, diffusing aggregative particles (Aktar et al., 2019; Krobitsch and Lindquist, 2000; Muchowski et al., 2000). We have found that mutant Htt-GFP in *S. cerevisiae* can diffuse throughout the IB, and that mHtt-GFP is released from the IB, indicating that the contents exchange with the cytoplasm, indicating that the IB has liquid-or gel-like properties (Aktar et al., 2019).

In addition to the experimental evidence indicating that material inside the inclusion body is diffusible, time-lapse imaging demonstrated that inclusions themselves are mobile (Aktar et al., 2019). Although it has been proposed that mHtt inclusions in yeast accumulate material primarily through active transport of the material to the inclusion (Hill et al., 2017; Kumar et al., 2016; Rothe et al., 2018; Wang et al., 2009), the movement of inclusions makes it difficult to envision how such a mechanism could be achieved. Furthermore, the smallest detectable particles of mHtt move entirely randomly, with no contribution by active transport. Therefore, we have proposed that the inclusion grows as a result of the coalescence of small particles of aggregated material into a larger inclusion (Fig. 1) (Aktar et al., 2019).

**Figure 1.**
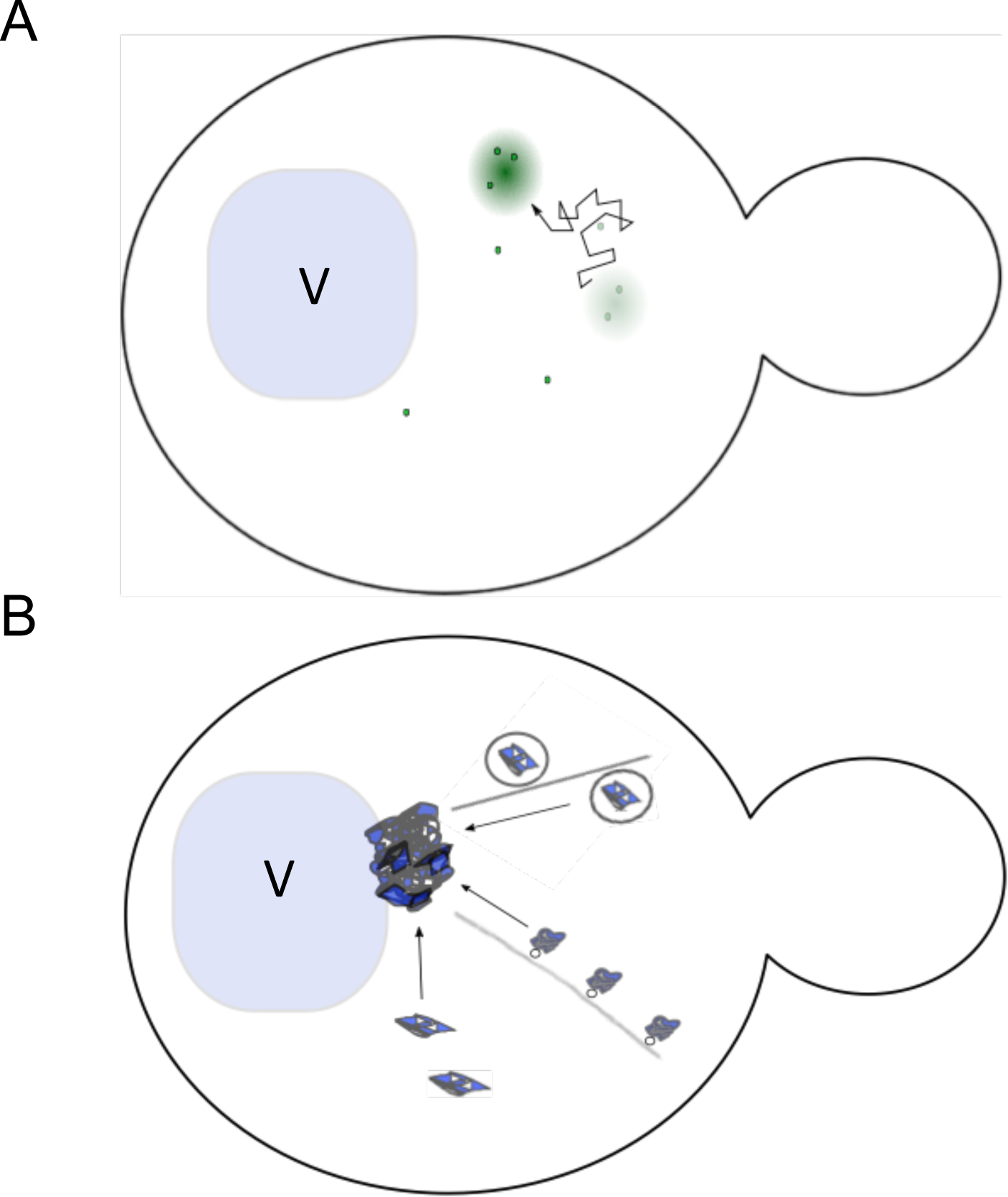
Two proposed models for growth of the mutant Htt inclusion in *S. cerevisiae*. **(A)** Model of mHtt inclusion as mobile phase-separated body: small particles move randomly in cytoplasm, and are integrated into the inclusion through collision and coalescence. **(B)** Model of mHtt inclusion as an IPOD: the IPOD is described as docked at the vacuolar membrane. Active transport along actin cables has been shown to be required for normal IPOD growth, and it has been proposed that material is transported in CVT vesicles or mediated by Myo2p. Transport by currently unknown mechanisms is also speculated to contribute to IPOD growth. V, vacuole.

In solution, or liquid-gel or liquid-liquid phase separations show a sharp concentration dependence (Banani et al., 2016; Ceballos et al., 2018; Trappe et al., 2001). The model of the mutant Htt IB as a phase-separated membraneless organelle that grows by collision and coalescence with small particles of unfolded protein predicts that IB formation should show a concentration dependence. At very low cytoplasmic concentrations, IBs should diminish because the rate of loss would outstrip exceed the rate of addition from random collision. Additionally, if material is accumulated through collision with the inclusion surface, we would predict that the growth rate of the IB volume should increase with the surface area of the IB, and it should also be modulated by cytoplasmic concentration.

We examined the concentration-dependence of IB formation in vivo and found that there appears to is a threshold concentration below which IBs do not form. Conversely, when we used auxin-mediated degradation to reduce cytoplasmic mHtt-GFP to low levels, material was lost from IBs. We observed that there appears to be a small kernel of mHtt-GFP that persists more stably, with a rate of loss that is markedly lower. Surprisingly, we also found that the proteasome is not typically overwhelmed in cells that form inclusions, indicating that it has a similar capacity to degrade mHtt-GFP in cells with and without IBs

We conclude that rising concentrations of unfolded protein trigger the nucleation of an inclusion body by the cell. Inclusion growth rate is consistent with the model in which material is absorbed after collision of aggregative particles with the surface. These studies constitute a novel, detailed analysis of the concentration-dependence and growth of a phase-separated body in vivo, and suggest that the process is nucleated and occurs independently of the capacity of the ubiquitin proteasome system (UPS).

## Results

### Time required to initiate an IB is not strongly dependent on mHtt concentration

We measured the time required to initiate a mHtt-GFP IB and IB growth. Inclusions composed of mHtt-GFP are typically singular, and do not appear to fission or fuse. In order to confirm that we would be able to track individual inclusions over long time periods, we selectively imaged 11 atypical cells with two (in one case, three) mHtt(72Q)-GFP IBs over a cumulative total of 35.7 hours, imaging every 10 minutes. We did not observe fusion of IBs. Nor have we ever observed fission of IBs, neither when imaging at 10 minute intervals, nor when imaging at frame rates of approximately 30 ms (Aktar et al., 2019). Thus, we are able to follow individual IBs over several cell cycles.

In order to determine the relationship between the time required for a mHtt-GFP IB to form and the cytoplasmic mHtt concentration, we used cells expressing mHtt(72Q)-GFP from a low copy number (CEN) plasmid (p415), under the control of the GPD promoter. The p415 plasmid has been found to be present at typically 2-5 copies per cell in our parental strain, BY4741 (Karim et al., 2013). Variations in plasmid copy number result in variable cytoplasmic concentration of mHtt(72Q)-GFP protein. In order to ensure uniformity of culture and imaging conditions, we used cells from a single mid-log phase culture, grown under identical conditions but with variable cytoplasmic concentrations of mHtt-GFP, to measure IB formation and growth.

As mHtt-GFP is continuously synthesized over the imaging timecourse, an individual cell contains a collection of GFP molecules that have been exposed to a variable number of exposures. As a result, we have found that it is not possible to correct mHtt-GFP fluorescence intensities accurately for photobleaching over long timecourses, due to continuous synthesis of mHtt-GFP (Aktar et al., 2019). To minimize any effect of photobleaching in this experiment, we imaged each field of cells only once per hour.

Forty individual cells growing on an agar pad and expressing mHtt-GFP were imaged every hour over a period of 5-6 hours. The original cells continued to grow and divide during the entire imaging period (Fig. S1), with a doubling time of 130 ± 7 minutes (mean ± SEM, n=27 initial cells), somewhat longer than the doubling time for the same cells in liquid culture with shaking (106 ± 2 minutes, n=10 cultures).

We tracked both cytoplasmic levels of mHtt-GFP and the formation of inclusion bodies (IB) in a total of 152 cells: the 42 initial cells and 110 of their progeny. Of those, 13 cells had an inclusion at the beginning of the experiment and 80 formed inclusions during the timecourse, while 59 cells did not form an inclusion during the period of our observations. Fig. 2 shows a timecourse of a typical cell dividing over 6 hours to give rise to 6 additional daughter cells.

**Figure 2.**
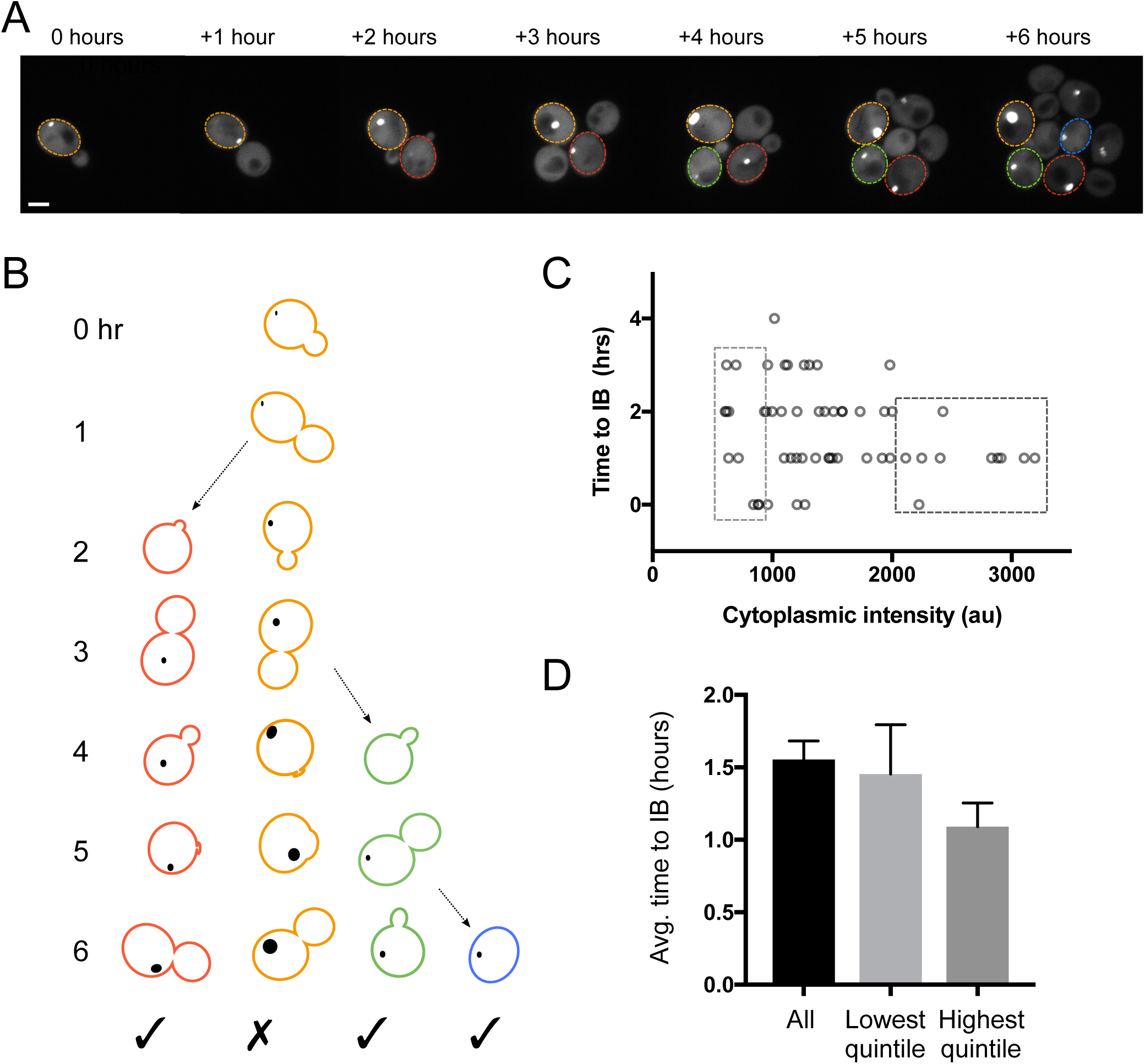
The time to formation of IBs is not strongly correlated with cytoplasmic intensity. **(A)** A typical field of cells imaged over a 6 hour timecourse is shown; the maximum projection includes those sections containing IBs. Outlines indicate the cells shown in (B). The contrast has been set to identical parameters for all images. Scale bar, 2 μm. **(B)** A diagram illustrating the criteria for inclusion in this analysis, showing the original cell in (A) and three cells born during the experiment. For the analysis in (C) and (D), measurements were only taken on cells born during the experiment that formed a new IB (indicated with a check). **(C)** Cytoplasmic intensity vs. the time after the birth of the cell that an IB first became visible. There appears to be a mild correlation between early birth and high intensity, but it is not strong enough to be significant (n_XY_= 59, Pearson’s *r =* −0.22, p = 0.11). Boxes indicate IBs that form in cells with cytoplasmic intensities in the lowest and highest quintile. **(D)** The average time to form an IB is shown for all cells, and for cells with cytoplasmic intensities in the lowest and highest quintile (mean ± SEM, n=56, 11, 11). The difference between time to IB formation is not significant (Kruskal-Wallis test across all 3 categories, p = 0.32; Mann Whitney test between all IBs and the highest quintile, two-tailed distribution, p = 0.12).

The cytoplasmic intensities of individual cells remained fairly constant over the course of the experiment (Fig. S2). In order to establish whether there is a relationship between cytoplasmic concentration of mHtt-GFP and the time to IB formation, we began by measuring the time from the birth of a cell to the formation of an IB. Our data set was restricted to cells that formed an IB and were born during the course of the experiment (Fig. 2B). Cells present at the beginning of the experiment were excluded because we could not know how long those cells existed prior to the onset of imaging. For cells that were born during the course of the experiment and formed an IB, the average time to IB formation was 1.6 ± 0.1 hours (mean ± SEM, n=56) (Fig. 2C, D).

Although there appears to be a weak inverse relationship between cytoplasmic concentration and the average time to form an IB in those cells that do form IBs, the correlation was not significant (n=56 XY pairs, p=0.11). If one considers the cells with cytoplasmic mHtt intensity in the lowest quintile, the average time to IB formation was 1.5 ± 0.4 hours (n=11), while the average time to IB formation was 1.1 ± 0.1 hours (n=11) for cells with cytoplasmic intensity in the highest quintile (Fig. 2D).

### Cells require a threshold cytoplasmic mHtt-GFP concentration to form an IB

Is there a concentration threshold required for inclusions to form? We monitored the intensity of cytoplasmic mHtt-GFP over the timecourse. While IBs formed in a wide range of cytoplasmic mHtt concentrations, cells that failed to form an inclusion often had very low cytoplasmic mHtt concentrations. Of inclusion-forming cells, 81% formed inclusions within 2 hours of the birth of the cell. Nine of the eleven cells in the lowest quintile of cytoplasmic concentration formed an inclusion in ≤ 2 hours. In our analysis of cells which did not form an IB, we considered only cells that were born over 2 hours before the end of the experiment, which were observed for 2 or more hours after the birth of the cell (Fig. 3A, B). Using this criterion, we identified 37 cells that did not form an inclusion during the course of the experiment, including 5 of the original cells present at the beginning of the imaging session, and 32 cells born during the experiment. These non-inclusion-forming cells were observed from 2–6 hrs, with a mean observation time of 3.6 ± 0.2 hrs (n=37, SEM).

**Figure 3.**
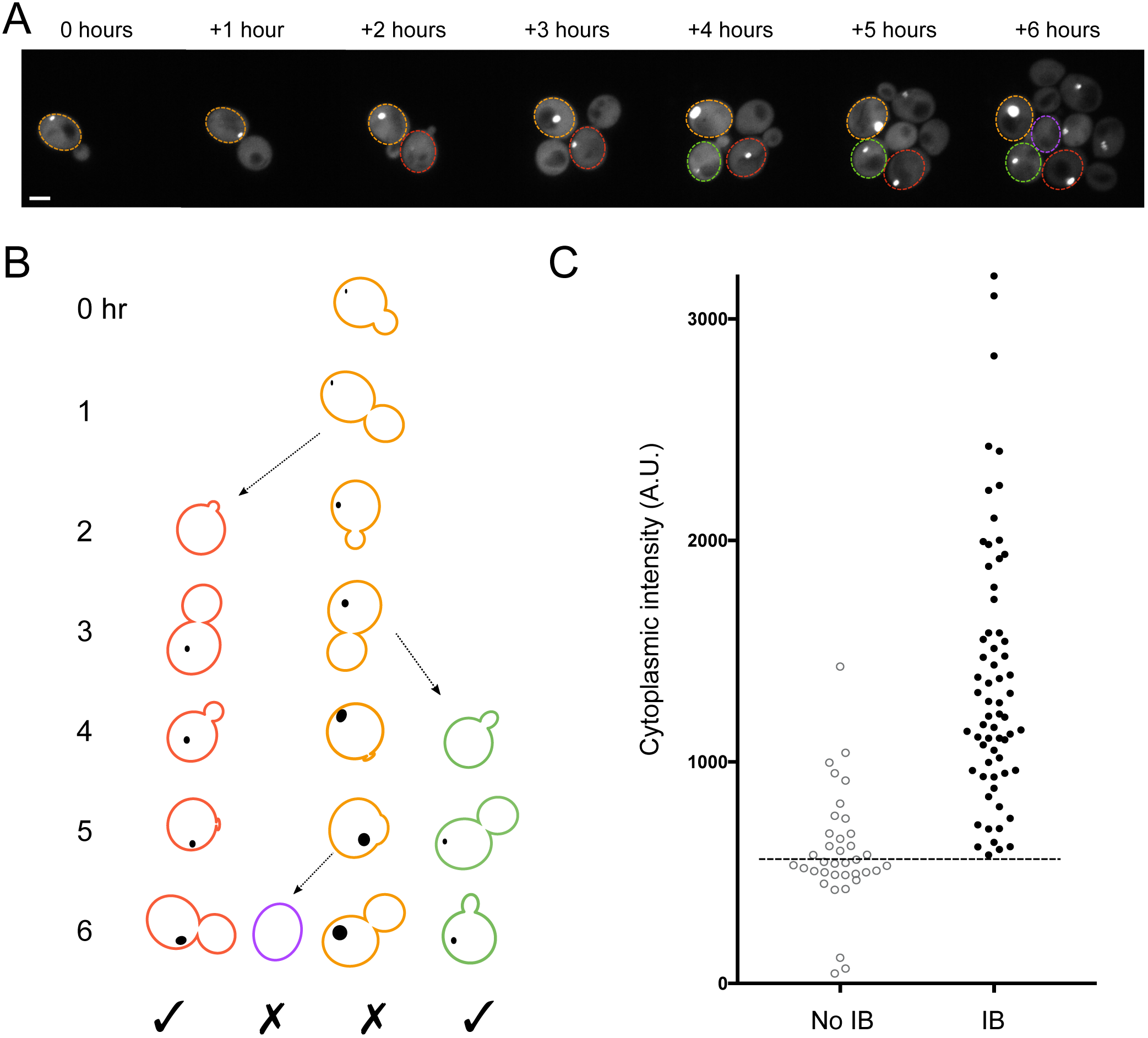
A threshold cytoplasmic concentration is required for IB formation. **(A)** The timecourse shown in Figure 2, but here the outlined cells indicate the four cells shown in (B). **(B)** In this analysis, we considered only cells that acquired an IB during the experiment and cells that did not acquire an IB but were born at least 3 hours before the end of the experiment. **(C)** Each data point reports the average cytoplasmic intensity of a cell that formed an IB (IB, n=65) or did not form an IB (no IB, n=37) during the course of the experiment. The intensities of the two groups are significantly different (p<0.0001, unpaired *t*-test, two-tailed distribution).

Our data are consistent with a concentration threshold of mHtt-GFP (Fig. 3C) for IB formation in growing cells. No cell with an average cytoplasmic concentration under 600 A.U. formed an inclusion during the period of our observations. Conversely, all but 2 of 102 cells with a cytoplasmic intensity >1000 formed inclusions. The average cytoplasmic intensity in IB-forming cells was 1485 ± 90 A.U (n=55, SEM), ranging from 605 – 3194.

### At low cytoplasmic mHtt-GFP concentrations, inclusion bodies shrink

The threshold cytoplasmic mHtt-GFP intensity for IB formation suggests that there is a concentration threshold required for either the initiation or the growth of an inclusion. Previous studies demonstrated that the IB is a phase-separated, although not liquid, compartment (Aktar et al., 2019). Phase separation is concentration-dependent and reversible *in vitro*. Thus, we next asked what would happen if the cytoplasmic concentration of mHtt-GFP were to drop *in vivo*? We used auxin-inducible degradation to drive down cytoplasmic mHtt levels and observe the effect on IB size. Auxin-inducible degradation was conferred by fusing the degron sequence (IAA^71-114^) to a target protein together with co-expression of the plant E3 ubiquitin ligase Tir1. Tir1 binds to the degron sequence in the presence of the plant hormone, auxin (1-naphthalene acetic acid (NAA)), resulting in the ubiquitination of the degron and degradation of the target protein (Nishimura et al., 2009; Papagiannakis et al., 2017).

Cells expressing Tir1 and mHtt(72Q)-degron-GFP were imaged every 15 minutes for 4 hours by spinning-disk confocal microscopy. The addition of auxin caused a substantial drop in cytoplasmic intensities of cells expressing mHtt-degron-GFP (Fig, 4A). As a control, we treated cells with the identical volume of vehicle (95% ethanol), which had no effect on cytoplasmic intensity (Fig. S3).

To study the fate of IBs during inducible degradation of mHtt, we identified cells containing IBs of moderate to large size (maximum diameters 0.6 −1.0 μm) that also showed a decrease in cytoplasmic intensity to background by about the 2^nd^ hour of imaging. The integrated density of the IB was measured over the 4-hr timecourse. As we have previously discussed empirical correction of the cytoplasmic intensity for photobleaching will substantially overcorrect as new material is continuously synthesized during the timecourse (Aktar et al., 2019). However, once the cytoplasmic concentration of mHtt-degron-GFP is very low, new material can no longer be added to the IB in significant quantities. Thereafter, the collection of molecules in the IB can be accurately corrected for photobleaching. Therefore, we have corrected all intensity data for photobleaching, but note that the corrected intensities prior to the loss of cytoplasmic fluorescence may be slightly inflated. However, we are most interested in the period after cytoplasmic levels dropped close to backgrounds, at which point correction for photobleaching is more accurate.

Inclusion body sizes vary widely from cell to cell at the beginning of the experiment. In order to compare the amount of material in IBs, we normalized the volume and integrated density of each inclusion to its maximum, and aligned the measurements at the point of maximum volume (Fig. 4B), or integrated density (ID) (Fig. S4A). Cytoplasmic intensities were also aligned to the point of maximum IB intensity, in order to determine the approximate relative cytoplasmic levels at which IB growth reverses.

**Figure 4.**
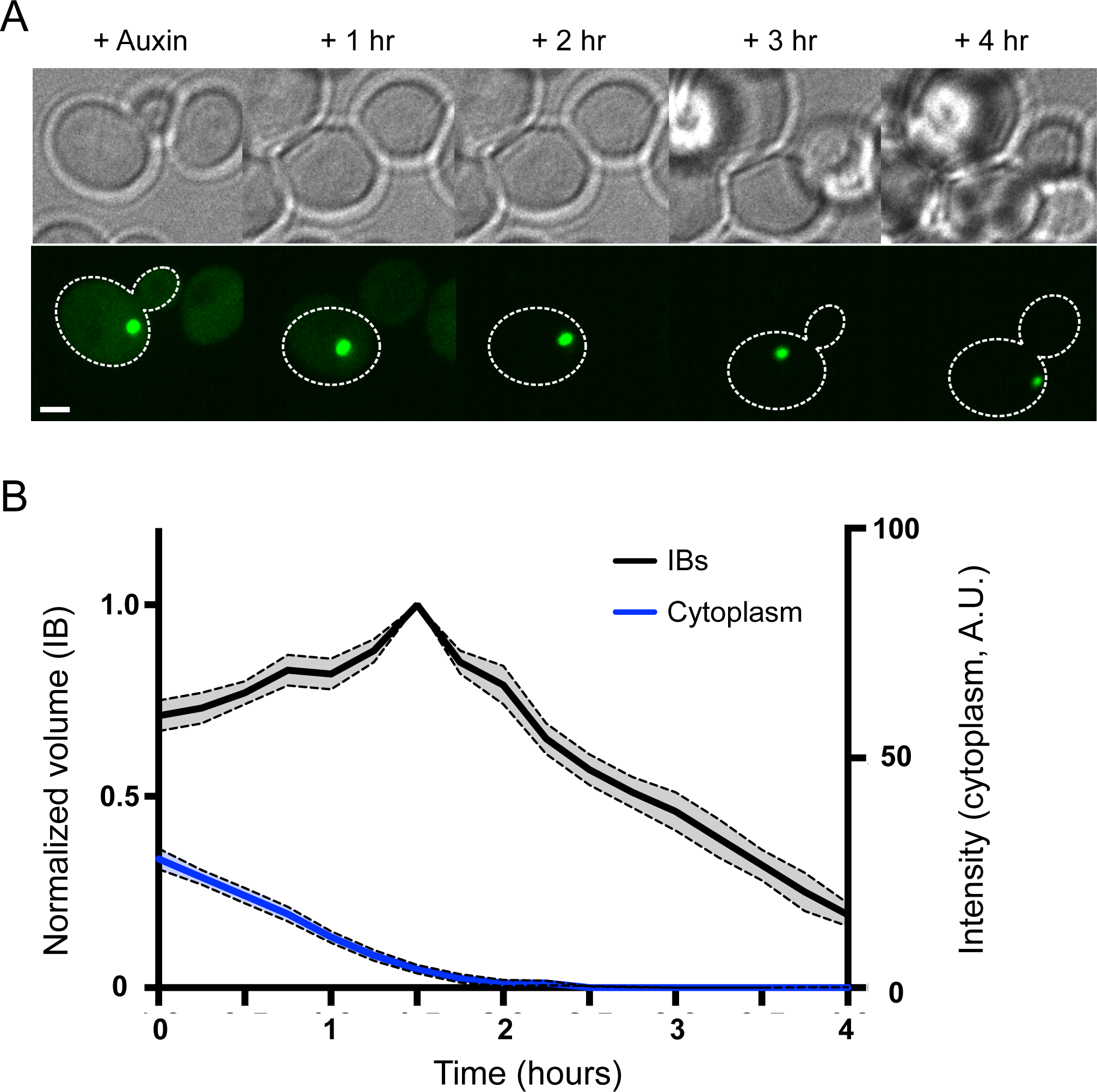
Mutant Htt inclusions shrink when cytoplasmic mHtt levels drop to low levels. **(A)** Time-lapse images of cells expressing Tir1 and mHtt-degron-GFP were acquired immediately after the addition of NAA, and every 15 minutes for 4 hours thereafter. Brightfield images (upper panels) and maximum-intensity projections of the GFP channel (lower panels) were taken shortly after the addition of auxin, then 1, 2, 3 and 4 hours later, illustrating loss of cytoplasmic GFP followed by decline in IB size. The displayed contrast of the fluorescence images is identical across the timecourse. The cell boundary is indicated by a dotted white line. Scale bar, 2 μm. **(B)** Quantitation of average IB volume (black) and cytoplasmic intensity (blue) over time. Volume traces were normalized to the maximum for each IB, and aligned in time so that the maximum intensity of each individual IB occurs at the same time point (n = 7-19 cells per timepoint; shaded regions indicate SEM).

Under the imaging conditions used in this experiment, cytoplasmic levels of mHtt-degron-GFP were 5-65 A.U. above background prior to the addition of auxin; the mean intensity of the cytoplasm was 22.5 ± 2.5 (n=25, SEM). Generally, inclusion bodies continued to grow as long as the cytoplasmic intensity of mHtt-degron-GFP remained at or above approximately 4-6 A.U., regardless of the presence of auxin. For all inclusions, at low cytoplasmic intensities, the volume peaked, then underwent a sharp decline when the cytoplasmic intensity of mHtt-degron-GFP fell close to background levels. Similar results were observed when integrated density was plotted over time (Fig. S4A).

In a small percentage of actively growing cells, a different form of inclusion body may be seen. We have called these inclusions ‘cluster-like’ (CLIs) because they have the appearance of a cluster of smaller inclusions clumped together: they are asymmetric in shape and their internal intensity is inhomogeneous (Aktar et al., 2019). We identified 4 cells containing CLIs and measured their response to loss of cytoplasmic mHtt-degron-GFP. Though a small sample (N=4), the response of CLIs was robustly different to the response of IBs: CLIs lost little of their contents. By either individual t-tests or two way ANOVA, the p-value for the difference in integrated density for IB’s vs. CLI’s in the last 5 time points was <0.001 (Fig. S4 A, B).

### The IB has a stable core

Moderate and large inclusions shrank relatively quickly, dropping to only 30% of their peak volume, or 22% of their peak integrated density, in 2 hours. In contrast, we observed that inclusions that were already small at the beginning of the experiment also lost material rapidly, but that a small, fluorescent core persisted until the end of the experiment, up to several hours. This core particle, or kernel, appeared to shrink at a much slower rate than the bulk of the inclusion (Fig. 5A).

**Figure 5.**
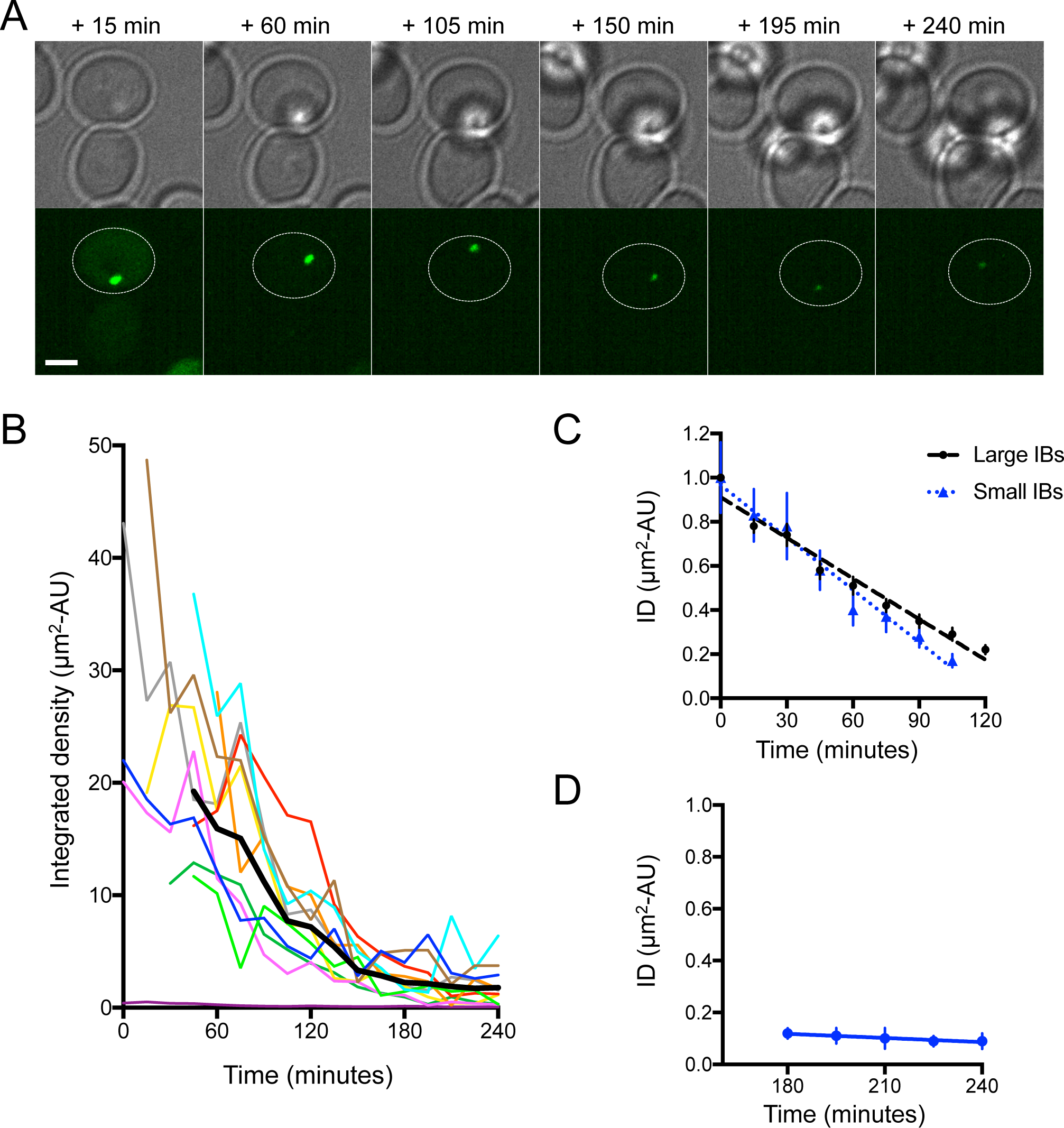
Biphasic loss of mHtt-GFP from the IB reveals a persistent core. (A) Brightfield (upper panels) and maximum-intensity projections of the GFP channel (lower panels) showing a cell containing a small mHtt(72Q)-degron-GFP inclusion after the addition of auxin. Time after addition of auxin is indicated above the images. The cell containing the IB is outlined. Scale bar, 2 μm. (B) Integrated density was plotted against time for 11 individual small IBs. The thick black line shows the average integrated intensity over time. IB intensities were corrected for bleaching. (C) The change in ID of small inclusions shown in (B) is shown for moderate and large (black) and small (blue) inclusions. The rate of decrease in ID is 37%/hr for large IBs (R^2^ = 0.97), and 47%/hr for small IBs (R^2^ = 0.97). Values are corrected for bleaching and normalized to initial IB intensity. (D) The rate of decrease in ID of small IBs during the last 75 min of the timecourse is shown (3%/hr, R^2^ = 0.94).

We tracked the core particles of 11 small IBs whose cytoplasm reached background levels ≤ 2hr after the addition of auxin. In order to compare the loss of material from persistent core particles with the loss of material from moderate to large inclusions, we plotted integrated density, as the apparent size of the IB core particles was well below our resolution. Of the 11 small IBs, none completely disappeared by the end of the timecourse, regardless of initial size (Fig. 5B). The initial loss rate was similar to that in moderate and large IBs (Fig. 5C). In contrast, at later stages, the average integrated density of the kernel was relatively steady (Fig. 5D).

### UPS has excess capacity in cells that form inclusions

Because inhibition of the proteasome can lead to formation of IBs, it has been proposed that IBs form in response to overload of the proteasome (Theodoraki et al., 2012; Tyedmers et al., 2010; Waelter et al., 2001); that is, the failure of one pathway of removal may lead to the formation of large aggregates of unfolded protein. Alternatively, blockage of the proteasome could allow cytoplasmic levels of unfolded protein to rise over a threshold required to trigger formation of an inclusion body. We were interested in ascertaining whether the proteasome was overwhelmed by unfolded protein in cells that had formed IBs or whether it still had excess capacity.

Consistent with our previous data, the cytoplasmic mHtt-GFP intensity appeared, on average, higher in cells with inclusions (Fig. S5). In a random selection of cells with and without inclusions, and subjected to auxin-induced mHtt degradation, we observe that cytoplasmic mHtt-degron-GFP levels decrease significantly over the course of the experiment in almost all cells: cytoplasmic mHtt-GFP levels fell in 100% of cells without IBs (n=86), and in 98% of the cells with IBs (n=43). In cells with an inclusion, the cytoplasmic signal remained visible for a longer period, but it dropped over the course of the experiment. However, we reasoned that the longer duration of detectability may have be the result of higher initial levels.

In order to compare the rate of auxin-induced cytoplasmic mHtt-GFP loss in cells with and without IBs, we selected cells with initial cytoplasmic levels between 5-45 A.U. that contained IBs, and compared them to cells with similar initial cytoplasmic levels, but no IBs, from the same imaging fields. Of 12 cells with moderate-to-large IBs, one did not significantly lose cytoplasmic intensity; the other 11 lost cytoplasmic mHtt-degron-GFP (Fig. 6A, B). The rate of mHtt-GFP loss from the cytoplasm in both cells with and without IBs was very similar: although the decrease in cytoplasmic intensity in cells with an IB was slightly slower, the difference was not significant (Fig. 6C). Since auxin-inducible degradation is mediated by the ubiquitin-proteasome system (UPS), these results are consistent with the supposition that the presence of an IB is not associated with lower UPS capacity.

**Figure 6.**
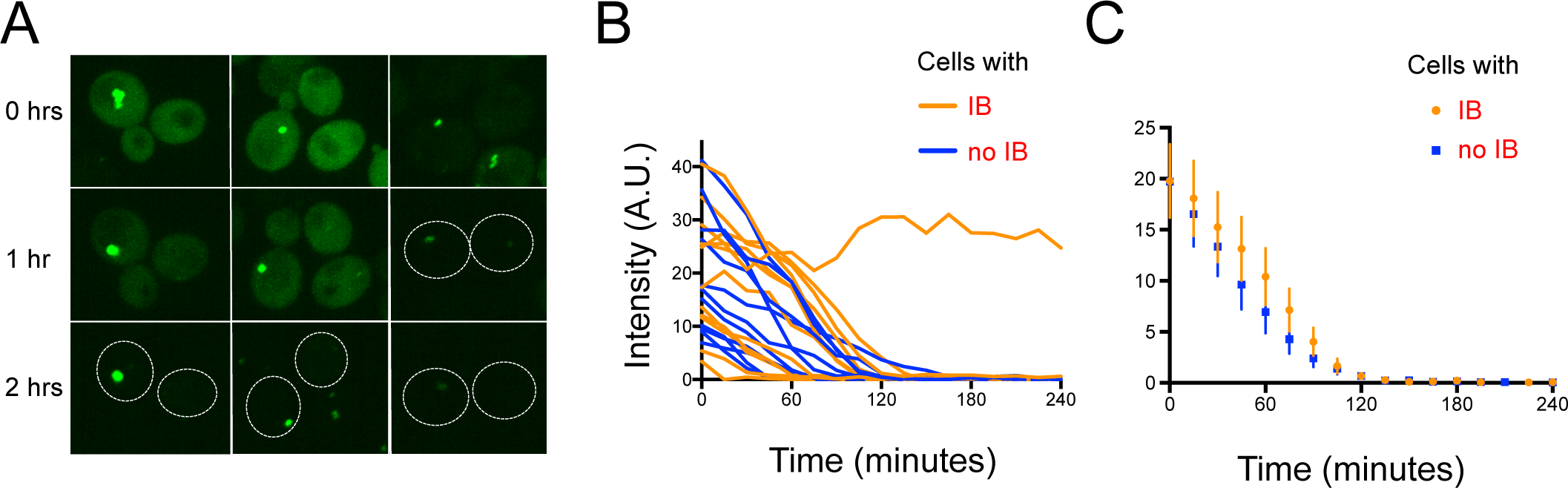
The proteasome has a similar capacity to acutely degrade cytoplasmic mHtt-GFP in cells with and without IBs. **(A)** Time-lapse images of cells expressing mHtt-degron-GFP were acquired every 15 minutes for 4 hours after the addition of auxin. Maximum-intensity projection of 3 pairs of cells with similar cytoplasmic levels is shown; in the cell shown in the upper left image, the inclusion was moving while the z-series was taken. Numbers indicate elapsed time. A confocal and brightfield image of the same field of cells is shown in Supp. Figure S5. Scale bar, 2 μm. **(B)** The cytoplasmic intensity of a random selection of cells with initial cytoplasm intensities between 5-45 AU with or without IBs, showing the loss of cytoplasmic mHtt-degron-GFP after the addition of auxin at time 0. **(C)** The average cytoplasmic intensities of the cells shown in (B) Error bars represent SEM; n=11.

### IB growth rate increases with surface area

Two fundamentally different models of inclusion growth in yeast have been proposed. Mutant Htt is most commonly described as accumulating in an insoluble protein deposit (IPOD), which has been proposed to receive a substantial fraction of new material through active transport (Fig. 1B; (Hill et al., 2017; Kumar et al., 2016; Rothe et al., 2018; Wang et al., 2009). However, the mHtt IB is mobile, making it difficult to transport material to it via existing cytoskeletal networks. Also, the movement of visible small particles is not directional. Therefore, we have proposed that the mHtt inclusion incorporates material through random collisions and coalescence.

Regardless of mechanism, we expect the growth rate of the inclusion to be proportional to the concentration of unfolded, aggregated mHtt, which, in turn, we expect should be proportional to the cytoplasmic concentration of mHtt. However, the mechanism of growth is predicted to affect IB growth kinetics. A model (Fig. 1A) in which growth occurs through incorporation of small particles onto the surface predicts that the rate of growth will increase with surface area, but may be modulated by the formation of a concentration gradient of diffusing particles. In contrast, the growth of IPODs has been shown to require functioning active transport systems; if material is actively transported to the inclusion, it suggests that growth may be dependent on the carrying capacity of the transport systems and independent of inclusion size. In the most extreme case, if transport is limiting, the transport model predicts a constant rate of growth over time as long as the cytoplasmic concentration of aggregates remains constant.

If aggregative particles are incorporated into the IB through collision, followed by incorporation of the particles, growth rate depends on the surface area of the IB and the speed with which particles are incorporated into it. The volume grows faster as the surface area increases because the frequency of productive collisions with small aggregative species increases with surface area. As cytoplasmic mHtt-GFP has been observed to be continuously replenished by new protein synthesis (Aktar et al., 2019), and to remain relatively constant with time (Fig. S2), it would be reasonable to predict that the concentration of mHtt particles will also be relatively constant over time.

Using those assumptions, there are two possible outcomes: if a stable concentration gradient of mHtt particles develops around the IB, we would expect growth to be limited by diffusion. In this case, we would predict that the surface area will grow linearly with time:

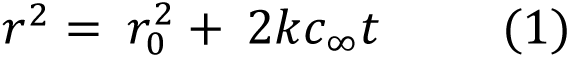

where *c*_∞_ is the bulk cytoplasmic concentration of mHtt particles, and *k* is the reaction constant, and *r* is the radius of the IB (see Methods).

However, if the rate of incorporation into the inclusion is slow and diffusion is sufficiently rapid that a concentration gradient of particles around the IB does not form, the concentration at the IB surface is the same as the bulk cytoplasmic concentration, and then we would predict that for any particular concentration of mHtt particles, the rate of growth of the radius will be constant over time:

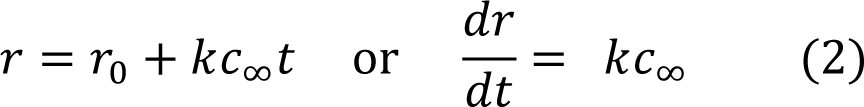

In the case of active transport, if the amount of material received by the IB is dependent on, and limited by, the volume of material transported to the IB, then the rate of change in volume over time can be expressed as:

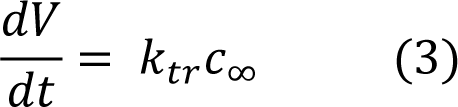

Here the identity of the aggregative species has not been defined and is not necessarily related to the visible diffusing mHtt particles. The rate constant is defined by the carrying capacity of the transport mechanisms.

In order to evaluate models of inclusion formation growth, we measured the growth of inclusions over time. Using the same long-term imaging data set that we used to relate cytoplasmic intensity to the formation of inclusions, we measured the growth rate of inclusions. We restricted our data set to inclusions that were > 0.3 μm in diameter and were followed for 4 or more hours (n=25 IBs). The initial diameter of these IBs was 0.48 ± 0.14 μm (mean ± SD). For each IB, we measured cross-sectional area in the optical section with the greatest maximum intensity in order to be as close to the center of the IB as possible. We used the cross-sectional area to estimate the volume of these ovoid structures at each time point.

A graph of all 25 IBs tracked for 4-6 hours shows that the growth rate increased with increasing average cytoplasmic levels of mHtt (Fig. 7A). For example, comparing several inclusions with similarly large size at the beginning of imaging (≈0.18-0.30 μm^3^), it is clear that inclusions in cells with high cytoplasmic intensity (1600-1700) grew significantly faster than those in cells with low cytoplasmic intensity (860-1270). These data confirmed our expectation that the amount of unfolded mHtt-GFP taken up by the IB is proportional to the overall cytoplasmic concentration of mHtt-GFP.

**Figure 7.**
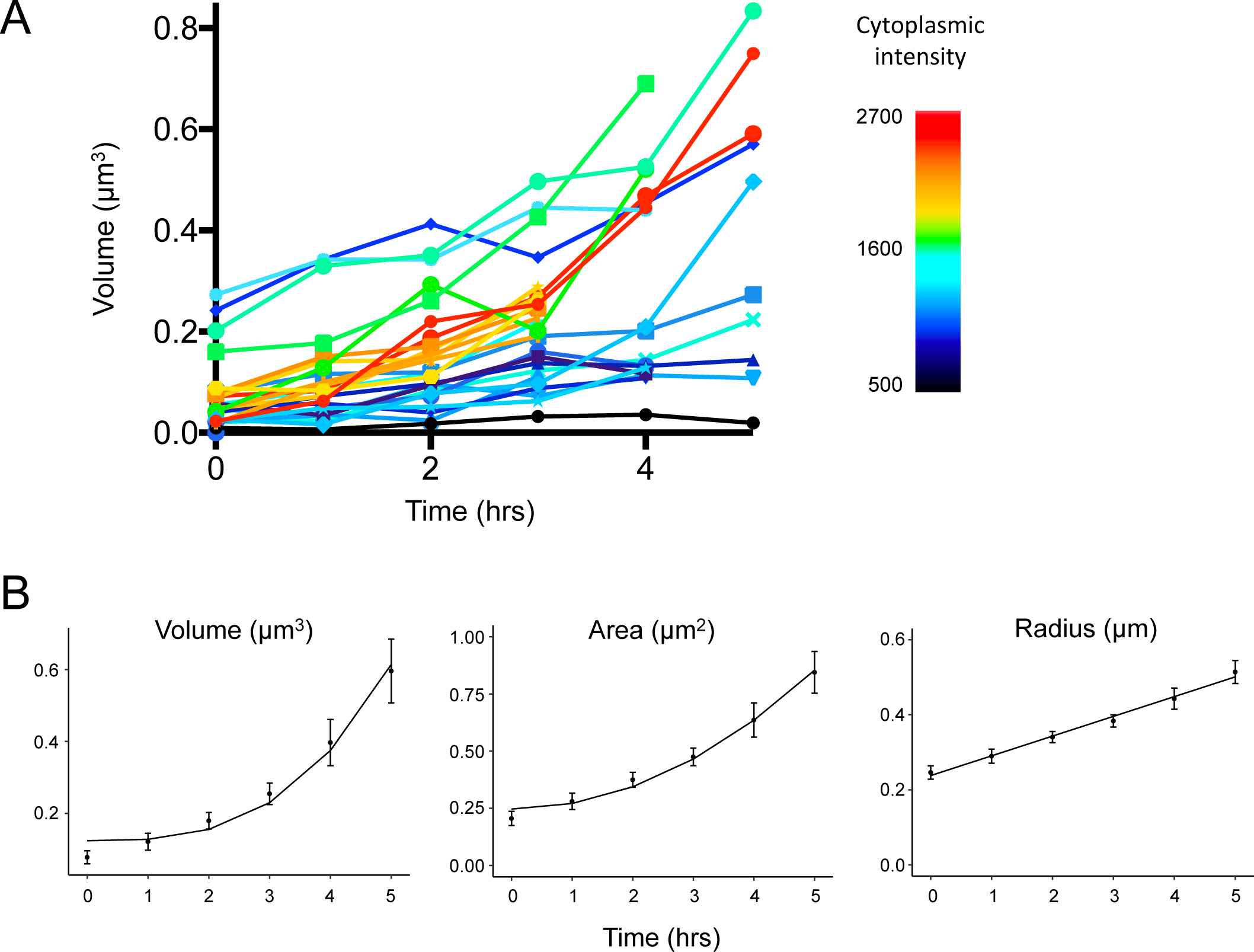
IB growth rate increases with cytoplasmic mHtt concentration and is proportional to IB surface area. **(A)** The measured volume of 25 IBs over 3-5 hours. The growth curves were aligned to time of first observation of each IB. The color of each IB trace indicates the average cytoplasmic intensity of the cell containing the IB (bar, right). **(B)** Average IB volume, area and radius are plotted over time for the 16 IBs in cells with cytoplasmic intensities >1100 (error bars indicate SEM). Fits of the growth of volume with *t*^*3*^ (R^2^ = 0.97), area with *t*^*2*^ (R^2^ = 0.98) and radius with *t* (R^2^ = 0.99) are shown.

To reduce noise in our analysis of growth rate, we used the cross-sectional area to calculate the approximate volume and radius of each IB, and then averaged the volume, cross-sectional area, or radius of all IBs. We fitted growth of volume, area and radius with time as a straight line and found that the growth of the IB radius was the best fit for a straight line with time, with growth of area with time also well-fit, either in the subset of cells with moderate or high cytoplasmic intensities (n=16), or all cells (Fig. 7B; Fig S6). If we extend our analysis to all possible implied relationships between the geometries with time, linear growth of the radius with time produces the best overall fits (Table S1). These results are consistent with a model in which growth is limited neither by the amount of unfolded protein nor by the rate of transport, but rather growth accelerates proportional to the surface area of the inclusion itself.

Additionally, the linear relationship between growth and IB radius suggests that the bulk concentration of mHtt particles is stable with time for cells with inclusions in the observed size range.

### The concentration of mHtt particles is proportional to that of soluble mHtt

Cytoplasmic mHtt-GFP has been shown to be largely soluble, based on biochemical studies (Krobitsch and Lindquist, 2000; Muchowski et al., 2000). In order to better understand the kinetics of mHtt unfolding, particle formation and IB growth, we wanted to know the relationship between cytoplasmic, soluble mHtt concentration and the concentration of unfolded mHtt particles that can be incorporated into the IB.

Unfortunately, we have not been able to directly visualize the bulk concentration of mHtt particles while simultaneously monitoring IB growth. Imaging of mHtt-GFP particles is possible, but requires sustained excitation that significantly bleaches the cell (Aktar et al., 2019). Therefore, we were not able directly assay IB growth and mHtt particle concentration in a single experiment.

However, the linear relationship between radius and time permitted us to assess the relationship between cytoplasmic mHtt concentration, and the underlying concentration of mHtt particles that were added to the IB. The data shown in Fig. 7A, B is consistent with the relationship:

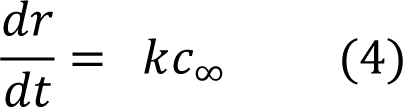

where *c*_∞_ is the bulk, cytoplasmic concentration of mHtt particles and *k* is the reaction constant for the incorporation of mHtt particles into the IB. We normalized and averaged the growth rate for 25 IBs coming from cells with cytoplasmic intensity ranging from approximately 500-2600 A.U. By fitting the data for individual IBs to a straight line, we obtained a slope equal to *k* with intercept at *r*_0_.

For each IB, we plotted radius vs time and fit the data with a straight line. Although these data are somewhat noisy, the coefficient of determination, R^2^, was ≥ 0.85 for 19 out of 25 IBs (Table S2). The slope of the line was calculated, giving us the value for kc_∞_ for IBs in individual cells.

We then plotted the value of kc_∞_ vs the cytoplasmic fluorescence, which is proportional to the concentration of soluble mHtt, for the cell containing that IB (Fig. 8). The relationship appears linear. Values for slope range from 0.01–0.08 μm/hour, and the root-mean-square error for the fitted line was 0.009. These data suggest that the concentration of mHtt particles that are competent to fuse with the inclusion is directly proportional to the concentration of the soluble, cytoplasmic pool of mHtt-GFP within this range.

**Figure 8.**
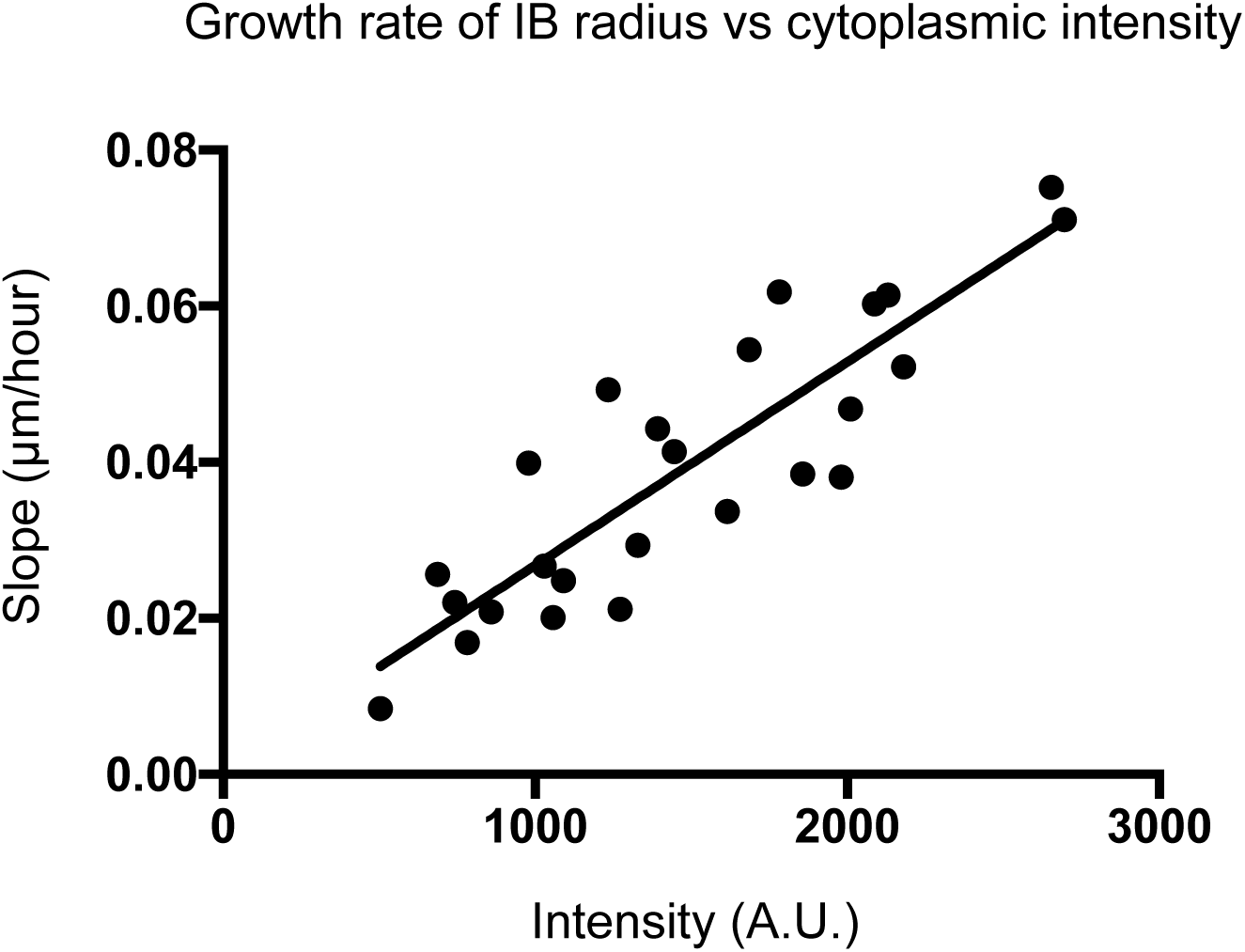
The concentration of mHtt particles is directly proportional to the concentration of soluble mHtt. The growth rate of the radius of individual IBs (slope of best-fit line) is plotted vs the intensity of the cytoplasm in which the IB was found. The distribution is fitted with a line (R^2^ = 0.77).

## Discussion

Our experiments provide insights into the mechanisms of IB initiation and growth, and allow us to create and test models of the dynamic partitioning of mHtt between a cytoplasmic phase and an IB phase. Expression of mHtt-GFP from a low-copy-number plasmid leads to natural variations in expression levels between cells of a population. We have further manipulated cellular mHtt concentrations using auxin-inducible degradation. Thus, we can use both natural and induced variation to probe the concentration dependence of inclusion formation and growth, and measure IB growth and shrinkage at diverse cytoplasmic mHtt-GFP levels.

Our results indicate that the formation of an IB is initiated, possibly by nucleation, at a threshold cytoplasmic concentration and regardless of proteasome capacity. The IB grows by collision and absorption of small particles of aggregated protein, and IB growth rate is dependent on the cytoplasmic mHtt-GFP concentration. IBs release material back into the cytoplasm, causing them to shrink when the concentration of cytoplasmic mHtt-GFP drops to very low levels.

Direct observation of cells with diverse cytoplasmic mHtt-GFP levels revealed a cytoplasmic steady-state threshold concentration below which cells never form IBs. Most (82%) of cells with mHtt-GFP concentrations over this threshold form inclusions, but not all do, suggesting that other factors play a role in IB formation. Conversely, in cells containing IBs where the cytoplasmic levels of mHtt-GFP have been acutely reduced by auxin-mediated degradation, the existing IB begins to shrink. Loss of material from the IB proceeds briskly until a small core particle, or kernel, remains.

The presence of a persistent kernel of mHtt after the bulk of the IB disappears suggests that there may be a non-phase-separated structure within the IB. It is possible that this represents a nucleation site. In solution, the presence of active centers can significantly lower the energy barrier to nucleation of a phase-separated compartment; seeding is used to speed the formation of phase-separated gels, for example (Rees et al., 2008; Shigekura et al., 2007). Nucleation is a mechanism by which the cell could control inclusion number and composition. Consistent with a nucleation model, we find that the time to form an IB is not strongly dependent on cytoplasmic mHtt-GFP concentration, unlike the growth of existing inclusions. This observation suggests that there is a regulated process that is initiated in cells with above-threshold levels of unfolded protein.

A prevalent model of IB formation states that inclusions form when the proteasome is overwhelmed, based on the increase in IB frequency seen when the ubiquitin-proteasome system (UPS) is inhibited (Theodoraki et al., 2012; Tyedmers et al., 2010; Waelter et al., 2001). Based on this model, we anticipated that cells with inclusions might not be able to carry out auxin-induced degradation because it acts through the UPS. However, we found that proteasomes can acutely degrade significant quantities of mHtt-degron-GFP in cells with and without IBs, suggesting that the UPS has excess capacity even when IBs have formed. Taken together, these observations suggest that IB formation is triggered by a regulated process when levels of unfolded protein reach a threshold, regardless of the capacity of the proteasomes.

Current models describe two fundamentally different mechanisms of growth for inclusions of mHtt-GFP (Fig. 1A, B) (Aktar et al., 2019; Hill et al., 2017; Kumar et al., 2016; Wang et al., 2009). In the phase separation model, the inclusion results from the diffusion and coalescence of small particles of aggregated material into a larger inclusion (Fig 1A). Alternatively, material may be actively transported to inclusions, as shown in Fig. 1B. In the active transport model, an IPOD is generally considered to be docked at the vacuolar membrane while a substantial fraction of new material is transported to the IB along actin cables, proposed to be mediated by Myo2p and/or Cvt vesicles (Hill et al., 2017; Kaganovich et al., 2008; Kumar et al., 2016; Rothe et al., 2018) In another version of the active transport model, the Sherman lab has proposed that the mHtt inclusion is an aggresome (Wang et al., 2009), located at the microtubule-organizing center and receiving material via retrograde transport along microtubules.

A key feature of the active transport models is that transport processes are limiting for inclusion growth. If transport is the limiting variable in inclusion growth, and the cytoplasmic concentration of material remains constant over the timescale of our experiment (consistent with our observations), then incorporation of new material into the IB should be linear with time.

Alternatively, the phase separation model predicts that the IB grows through integration of particles of unfolded protein captured by its surface. This proposal is consistent with the observed mobility of the IB and small particles, and would be characteristic of other phase-separated compartments observed in cells. A model in which Htt-GFP IBs incorporate small, diffusing particles of mHtt-GFP would also account for the finding that other Hsp104-dependent inclusions have been shown to contain subparticles of less soluble material (Bagriantsev and Liebman, 2004; Bagriantsev et al., 2006; Kryndushkin et al., 2003).

If the IB grows through collision with small particles of unfolded mHtt, the amount of new material taken up by the inclusion should increase with increasing surface area, as the growing surface will result in a greater number of collisions. However, the incorporation of particles into the IB may be limited either by the rate at which particles diffuse to the IB surface, or by the speed of the incorporation reaction. If diffusion of particles to the surface is limiting, we would expect the cross-sectional area to grow linearly with time (1). If diffusion is not limiting, then we would expect the IB radius to grow linearly with time (2).

Our measurements suggest that the inclusion radius does increase linearly over time. In this way, the growth of the IB may be compared to the growth of nanoparticles in solution. For the case where the surface reaction rather than diffusion of monomers is the limiting factor, then the change in radius with time has been described as

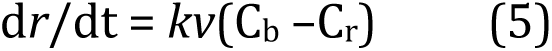

where *r* is the radius of the crystal, *k* describes the rate of the surface reaction, *v* is the molar volume of the monomer, C_b_ is the concentration of the monomer in bulk solution, and C_r_ is the solubility of the particle (Thanh et al., 2014). In our case, the cytoplasmic concentration of mHtt-GFP is relatively constant for each cell during the 5-6 hour timecourse. We assume, for moderate to large inclusions, that the surface concentration of mHtt-GFP particles is also constant, which we believe to be reasonable for a large body whose surface is not undergoing large changes in curvature (surface tension).

Lastly, we conclude that the concentration of mHtt particles that are that are competent to fuse with the IB is linearly proportional to the cytoplasmic, soluble mHtt-GFP concentration. The growth of the IB radius is constant with time

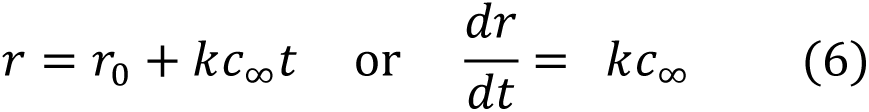

suggesting that the bulk concentration of mHtt particles, *c*_∞_, remains reasonably constant in our cell population. For each individual IB, we are able to calculate the value for *kc*_∞_ (6). When we plotted the value of *kc*_∞_. for each IB vs the cytoplasmic intensity of the cell containing that IB, we saw that *c*_∞_ increases linearly with the measured cytoplasmic intensity. Formation of mHtt-GFP particles is dependent on the dual-function aggregase/disaggregase Hsp104 (Aktar et al., 2019; Shorter and Lindquist, 2004; Weber-Ban et al., 1999). These findings suggest that substrate levels are not nearing saturation for Hsp104 in our cells.

Our imaging studies, consistent with earlier biochemical studies, suggest that the inclusion bodies are composed of discrete, insoluble particles that are absorbed into a larger phase-separated structure through collision, followed by coalescence. When the cytoplasmic concentration of mHtt-GFP is lowered, the inclusion releases material. Given that (1) increasing concentration has been shown to be the “switch” that flips Hsp104 from aggregation to disaggregation, and (2) the IB brings together Hsp104 and its substrate at high concentration in the inclusion, we suggest that formation of the inclusion body in yeast functions to collect unfolded protein and facilitate its disaggregation by Hsp104. In short, rather than a garbage dump for unfolded protein, inclusions are a recycling center.

## Materials and Methods

### Strains, plasmids and culture conditions

Yeast strains were grown using standard conditions and the appropriate glucose-based selective media at 30°C (Sherman, 2002). The lithium-acetate method was used for transformations; transformants were selected on the appropriate selective medium (Gietz and Schiestl, 2007). The doubling time of the mHtt(72Q)-GFP strain in liquid culture was determined using cultures grown in SC-Leu at 30°C with shaking. Overnight cultures were inoculated to OD_600_ = 0.03-0.05 and left to shake for 2 hours, after which cells were reproducibly in exponential growth phase. The cultures were sampled approximately every hour for 4-5 hours. Doubling time was calculated by plotting time vs. ln(OD_600_), fitting with a straight line (R^2^≥0.99), and dividing the natural log of 2 by the slope.

Plasmids used in this study are listed in Table S3. Mutant huntingtin expression plasmid pEB4 (mHtt(72Q)-GFP) was constructed as described previously (Aktar et al., 2019). The insertion of the IAA71-114 degron sequence into pEB4 between the Htt and GFP was made using standard PCR-based subcloning methods; Kapa Hifi (KapaBiosystems) or Phusion DNA polymerase (New England Biolabs) was used for PCR reactions requiring high fidelity. All plasmid DNA sequences derived from a PCR amplicon were sequenced and found to be free of mutations that change coding or known regulatory sequences. Integration of Tir1 into the HO locus was carried out using a linearized plasmid (Papagiannakis et al., 2017). Strains used in this study are listed in Table S4. The parental strain for the strains used in this study was BY4741 *MATa his3Δ1 leu2Δ0 met15Δ0 ura3Δ0*.

### Confocal image acquisition and data analysis for IB growth and cytoplasmic intensity studies

Cells were inoculated in selective medium and grown overnight at 30°C with shaking; cultures in mid-log phase the next day were selected for imaging study. When performing time-lapse experiments, cells carrying pEB4 were mounted on a 2% agar pad made with selective media, covered with a #1.5 coverslip, and sealed with valap to prevent drying. Fields of cells were selected in brightfield mode, and imaged every hour for 5-6 hours. Individual images from the same stack shown together are displayed with consistent contrast settings.

Cells were imaged using a 100x/1.45 CFI Plan Apo Lambda objective lens on a TiE2-PFS microscope (Nikon) equipped with a CSU-X1 spinning-disk unit (Yokogawa Electric, Tokyo, Japan), a Zyla sCMOS camera (Andor, Belfast, Northern Ireland) and OBIS LX 488 and LS 561 lasers (Coherent Inc, Santa Clara, CA).

Quantification was performed on unprocessed images using the Fiji distribution of ImageJ (Schindelin et al., 2012; Schneider et al., 2012). Using a script written by T.C.S. for ImageJ, the size and mean intensity of the inclusion body were measured in the optical section with the highest maximum intensity, assumed to be the section closest to the center of the inclusion. First, the mean cytoplasmic intensity was measured in a region excluding the vacuole and distant from the inclusion. After identifying the optical section with the highest maximum intensity, the inclusion was segmented using a threshold of 1.5x the mean cytoplasmic intensity. The Analyze Particles function was used to measure the area, position and mean, minimum and maximum intensity of the inclusion. Mean cytoplasmic intensity was also reported.

Previous work has established that mutant Htt(72Q)-GFP is synthesized and removed on a timescale that prevents correction for photobleaching in imaging studies conducted over a period of hours. At longer times, the population of molecules has been exposed to varying exposure times (Aktar et al., 2019). For this reason, excitations were widely spaced.

Estimates of IB movement on a short timescale (≤100 ms) were made as previously described (Aktar et al., 2019). In brief, time-lapse images were acquired at 30-32 fps and denoised with the Advanced Denoising function of NIS Elements Advanced Research software v.4.6. Particles were tracked using the SpotTracker plugin (Sage et al., 2005), and the average mean squared displacement was calculated from the particle coordinates determined by SpotTracker.

### Calculation of integrated density and size and line-fitting

Integrated density was calculated as the mean inclusion intensity multiplied by the area of the inclusion; mean inclusion intensity was corrected for background intensity. Inclusions are close to circular (Aktar et al., 2019) and were approximated as spherical for the purpose of calculating diameter, area and volume.

For each inclusion, the area through the plane of maximum intensity was taken as the maximum area for each inclusion. Radius was calculated as (area/π)^1/2^, and volume was calculated as 4/3 x area x radius (4/3πr^3^). Calculations were done in Excel (Microsoft Office), Prism (GraphPad, San Diego) and R (R Core Team, Vienna, Austria, 2020; https://www.R-project.org). Prism and R were used for graphing functions and statistics. Line fitting in Prism and R was used to determine the slope, standard error, 95% confidence interval and value for R^2^ of the best-fit line for IB radius vs time.

### Auxin treatment and image analysis

MatTek dishes (MatTek Corporation) were prepared by adding approximately 100 μL of a filter-sterilized 2 mg/ml solution of concanavalin A, allowing them to stand for 30 min at room-temperature, rinsing with distilled water and allowing them to dry overnight (Higuchi-Sanabria et al., 2016). One hundred microliters of an overnight culture of log-phase cells were added to MatTek dish and allowed to settle for 10 minutes before being rinsed gently with SC-leu. After rinsing, 200 μL of SC-leu was added to the dish. Fields of cells were selected in brightfield. Prior to the addition of auxin, two fields of cells were imaged 10 times as rapidly as possible to measure photobleaching, using the same settings that were used in the imaging experiment.

A fresh solution of naphthalene acetic acid (Sigma-Aldrich) was prepared to 60mM in 95% ethanol and diluted to 2.7mM in water. Twenty microliters of 2.7 mM naphthalene-acetic acid (NAA) was added to the center of the well, for a final concentration of 250 μM. Eight new fields of cells were imaged every 15 minutes for 4 hrs. For vehicle controls, 20 μL of 95% ethanol was added to the cells instead of NAA.

To correct for photobleaching, using the images collected before auxin treatment, five cells without an inclusion were selected and a region of interest was drawn to avoid the vacuole. The mean cytoplasmic intensity in cells was measured over 10 imaging cycles. The intensity measurements were normalized to the initial intensity, averaged and a line was fitted using an exponential decay function. The slope of the line and number of rounds of imaging were used to determine fraction of fluorescence loss due to photobleaching and correct mean intensity in auxin-treated cells.

Using the script described above, size and mean intensity of the IB in the optical section with highest intensity was determined. Intensities of IBs were corrected for photobleaching, as the population of IBs used in this study ceased incorporation of mHtt-GFP soon after the onset of imaging and therefore contained a stable or declining population of molecules that one could reasonably assume were subjected to the same, or very close to the same total number of excitations. Integrated densities and volumes were calculated as described above.

In our analysis of the loss of fluorescence in moderate to large IBs, a FIJI plugin was used to automate measurements of inclusion size. In the analysis of the IB core particle, however, the particle was identified manually and a region of interest was drawn around it. A threshold intensity of 1.1 above the background level using the Threshold function in ImageJ, and the Analyze particles function, were used to measure mean intensity and cross-sectional area.

### Inclusion growth rate calculations

#### Diffusion and collision-driven process

If we assume that the IB grows as a result of the incorporation of diffusing aggregative particles, we can calculate the mass balance on the IB as follows.

The mass *m* of the IB increases due to the flux of aggregative particles to the surface of the IB. We assume the origin of our coordinates to be at the center of the IB, taken as a sphere:

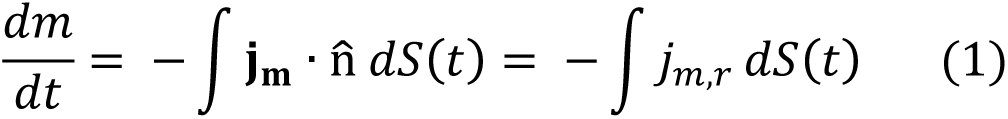

where *j*_*m,r*_ is the radial component of the mass flux **j**_**m**_, and 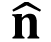 is the outward-facing unit normal from the surface of the IB. As it is evident that the IB grows, we can say that material is moving toward the origin, or 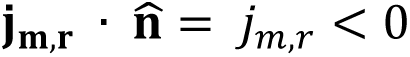. The mass is *m* = *ρV*, and assuming that the IB as a constant mass density, *ρ*, and spherical symmetry, this equation can be written for volume growth:

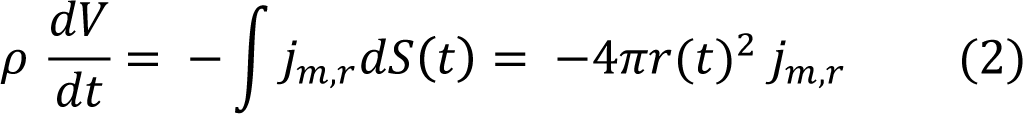

For the assumption of a spherical IB,

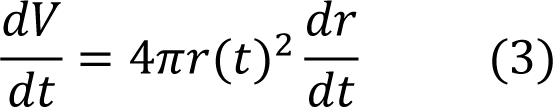

and (2) can be rewritten as

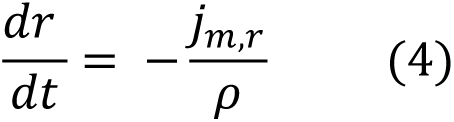

The flux *j*_*m,r*_ can be written in volumetric form *j*_*r*_ by using the density of the aggregative particles, i.e. *j*_*m,r*_ = *ρ*_*part*_, to yield

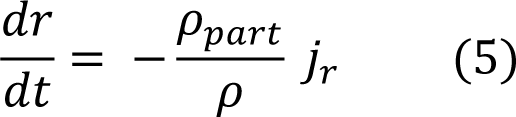

where it can be recalled that *ρ* = *ρ*_*IB*_. It is not clear that the density of the aggregative particles are the same as the density of the IB, which appears to have a heterogenous structure. However, if we take *ρ_part_/ ρ* as equal to a constant κ_den_, we can say

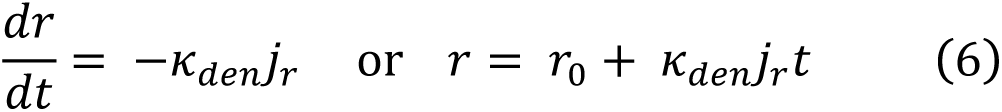

If we assume a steady state concentration develops around the IB, we can say that the concentration must satisfy the diffusion equation:

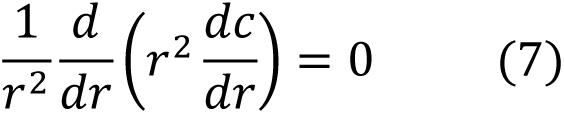

The boundary conditions are *c(R* → ∞) = *c*_∞_, and the surface flux at *R* = *r*, where *r* = *r(t)*:

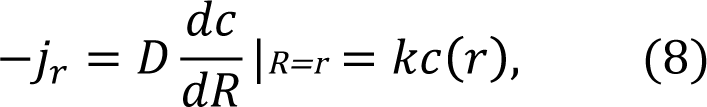

where *D* is the diffusivity of the aggregative particles in the cytoplasm, and *k* is the rate constant described above. The solution is *c* = *A+B/R* with *A* = *c*_∞_ and *B* is determined from the flux balance to yield

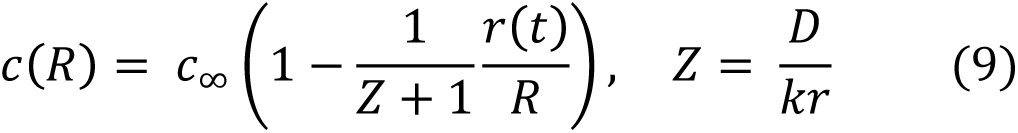

where *Z* may be considered as a dimensionless diffusivity, and is the inverse of the commonly used Damköhler number, Da = 1/ *Z* = *kr/D*.

For *Z* ≪ 1 (i.e., Da ≫ 1), the reaction rate is fast and limiting cytosol concentration as the the surface is approached vanishes, *c*(*r*(*t*)) → 0. In this case, we may rewrite (8) as

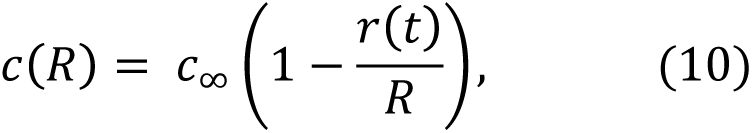

and the flux is

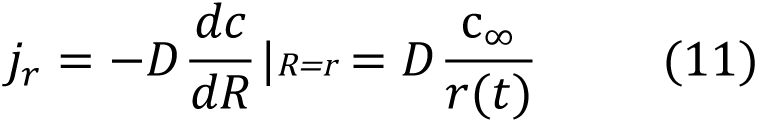

For fast reaction rates, we can say

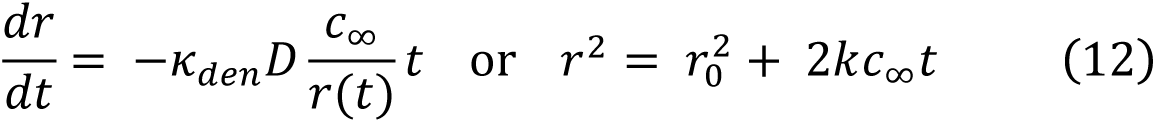

In this case, the surface area is predicted to grow linearly with time.

If the reaction rate is slow, *Z* ≫ 1 (or Da ≪ 1), and the combination of aggregate diffusion and movement of the IB has time to make the concentration field uniform (i.e. these factors prevent a stable concentration gradient from forming around the IB), then the concentration of particles at the IB surface, *c*(*r*(*t*)), will asymptotically approach the bulk concentration of particles in the cytosol, *c*_∞_. In this case, we may say that

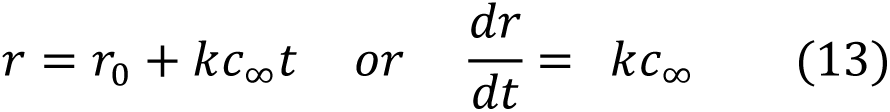

where *k* is the forward reaction constant for the integration of particles into the IB.

#### Transport-driven process

If we assume that material is transported to the IB by active transport systems, then the rate of material reaching the IB is dependent on, and limited by, the carrying capacity of the transport systems. We assume that the volume of material carried to the IB is constant with time, then we can say

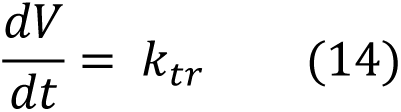

where *k*_*tr*_ is the rate constant for the transport systems delivering material to the IB in volume of material per unit time. If the capacity of the transport system is limiting, at some time after initiation of the IB, we can say

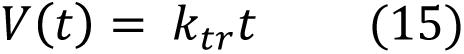

Substituting *4/3 πr*(*t*)*3* for volume, we can say

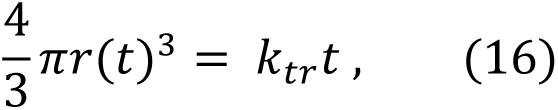

which can be rearranged to give:

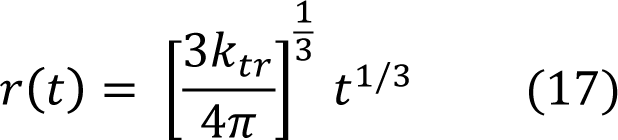

Therefore,

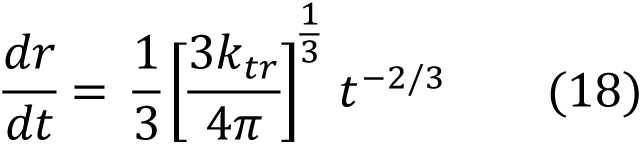

in the case where material is transported to the IB by transport systems with a capacity that is independent of the size of the IB.

## Acknowledgements

The authors thank Liza Pon, Istvan Boldogh, Janeska de Jonge and Enrique Garcia for their valuable advice and assistance with strain construction. We are grateful to Peter Lipke for his advice on the manuscript. We would also like to thank Pat Hooper for helpful discussion. This work was supported by grants from the National Institutes of Health to LE (1SC2GM116697). Images were collected on a Nikon Spinning Disk microscope acquired by Department of Defense Equipment Award W911NF-17-1-0516.

## Author contributions

L.E. conceived the ideas, performed the experiments and analyses, and wrote the manuscript. S.P. made strains and performed experiments. J.F.M derived analyses of growth. T.C.S. contributed to image analysis.

