## Supplementary Figures for "Threshold concentration and random collision determine the growth of the phase-separated huntingtin inclusion from a stable core"

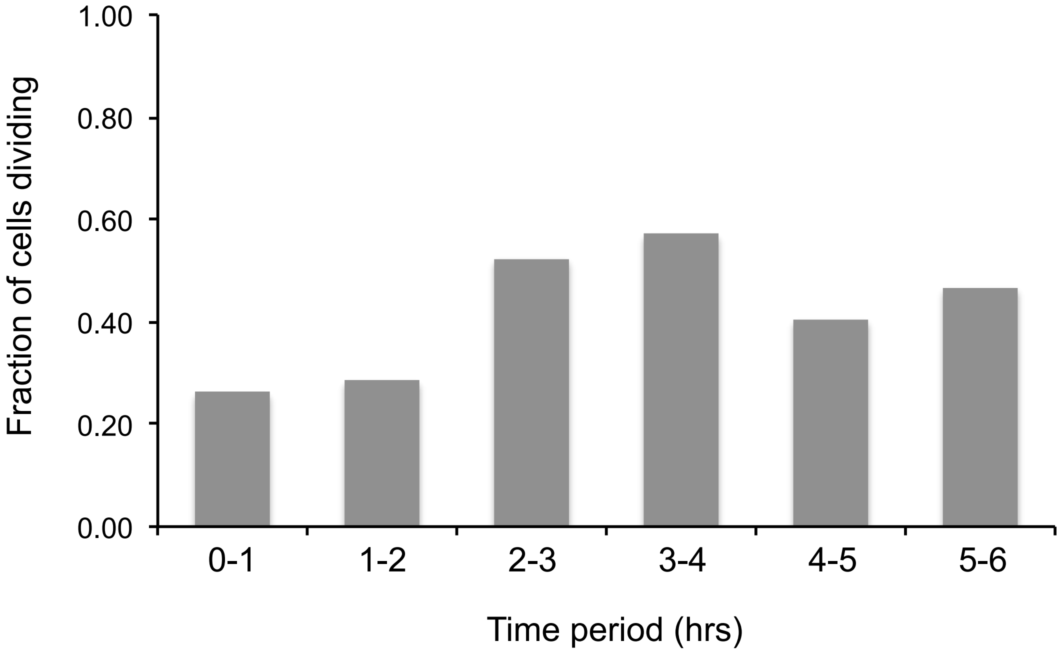


**Figure S1. Cells continue to divide throughout the imaging session.** For the 42 initial cells, the percentage of cells dividing within the indicated time period is shown.

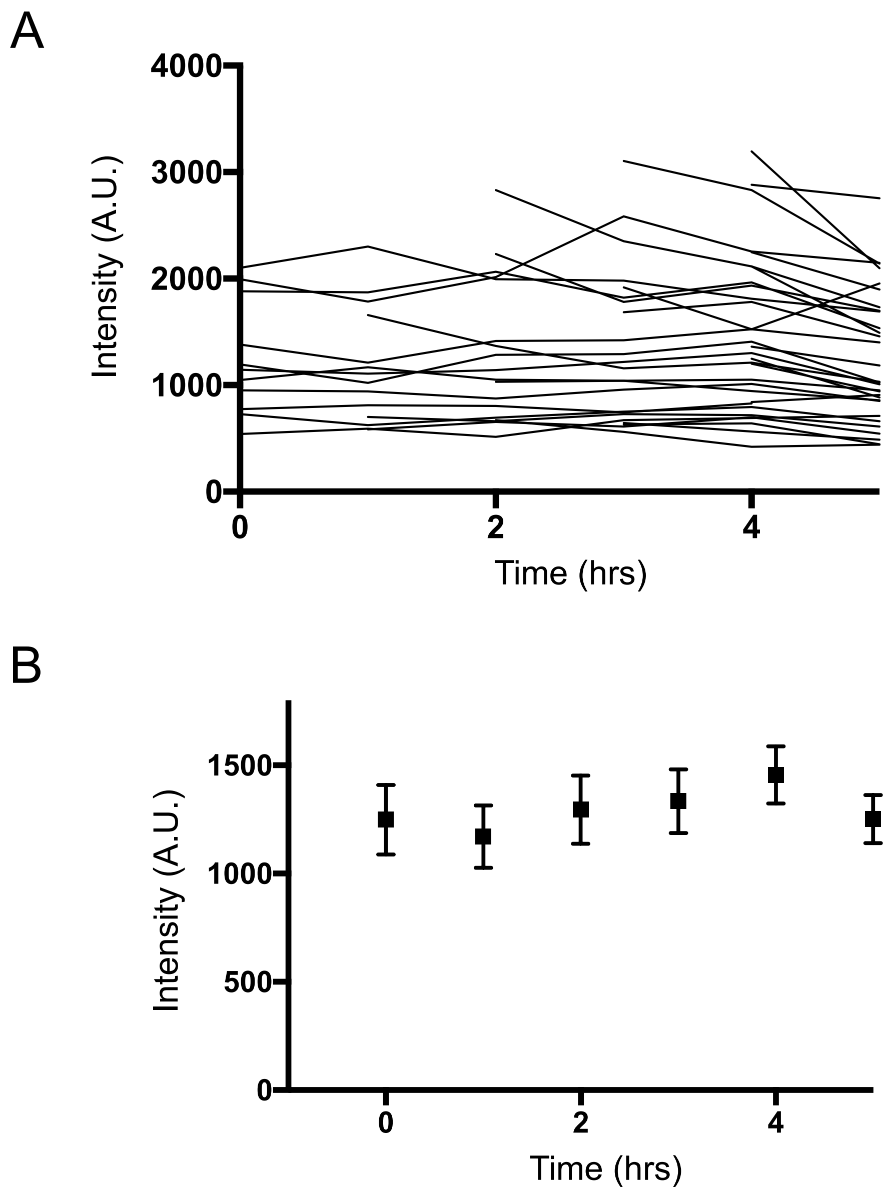


**Figure S2. Cytoplasmic mHtt-GFP intensity is stable over 5 hours of imaging. (A)** Cytoplasmic intensities over time for 31 randomly selected IB-forming cells are shown. Traces beginning after t = 0 represent cells that were born during the course of the experiment. **(B)** Mean intensity ± SEM is shown for the same 31 IB-forming cells.

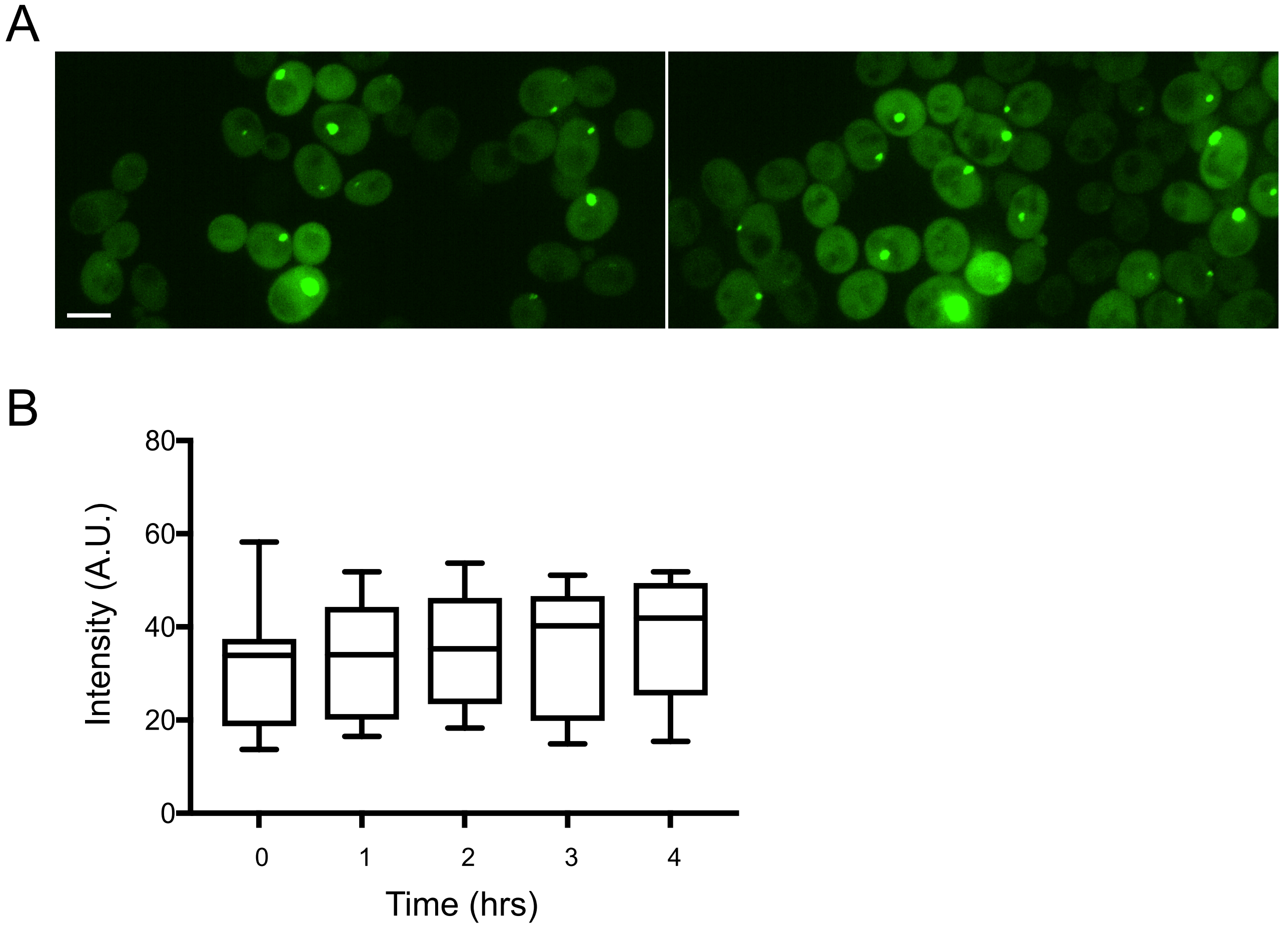


**Figure S3. Vehicle-treated cells expressing Tir1 and mHtt-degron-GFP show no change in cytoplasmic intensity. (A)** Cells grown identically to those in Fig. 4 were treated with the vehicle (95% ethanol) and imaged every 15 minutes for 4 hours. A representative field of cells at the beginning (0 hrs) and end (4 hrs) of the timecourse. Scale bar, 4 μm. **(B)** Average cytoplasmic intensity of 11 randomly sampled cells (whiskers indicate minimum and maximum values).

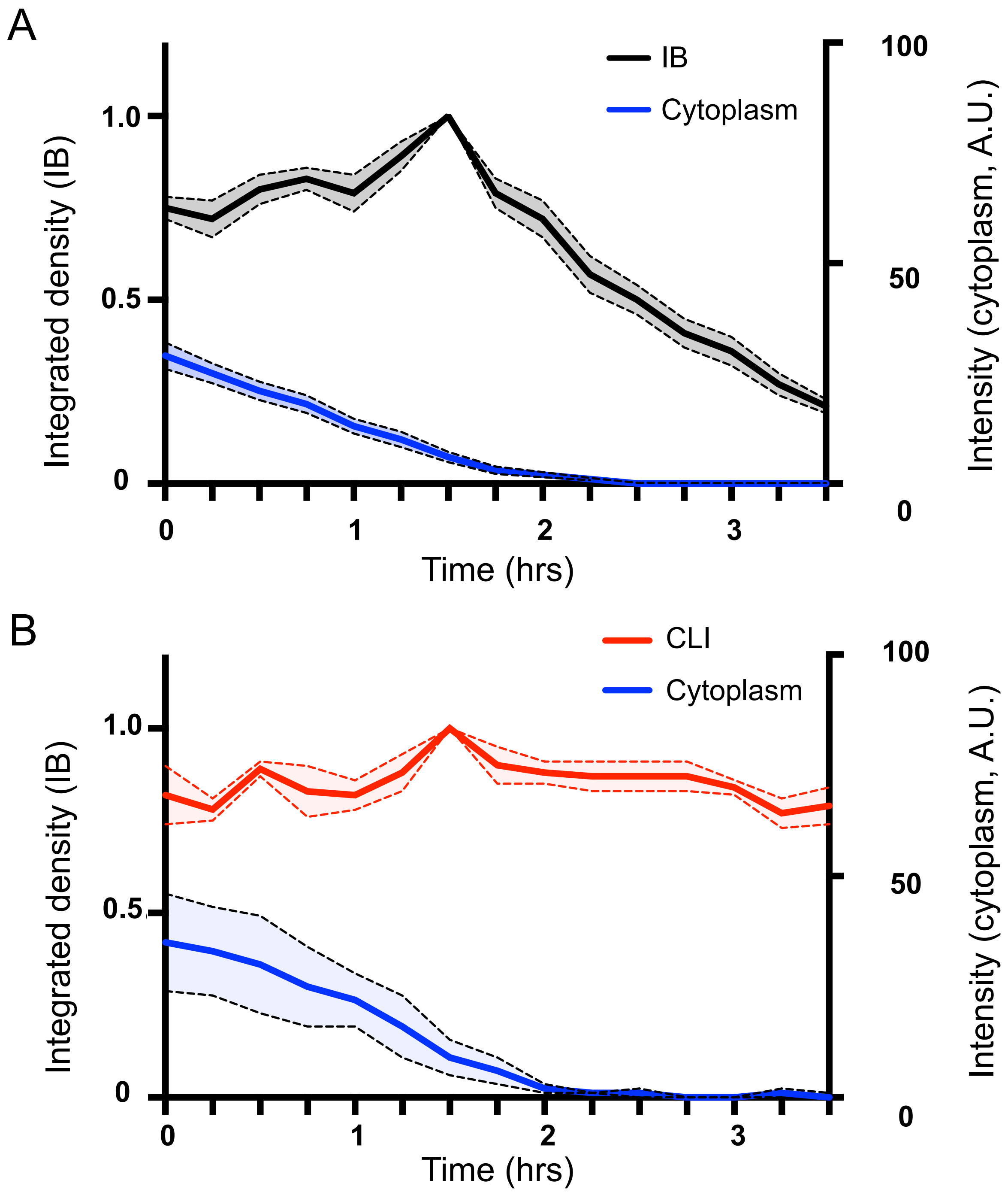


**Figure S4. Mutant Htt inclusion bodies, but cluster-like inclusions, decrease in integrated density when cytoplasmic mHtt levels drop to low levels.** (A) Normalized integrated density of IB (left axis) and cytoplasmic intensity (right axis) for 19 auxin-treated cells, showing mean (solid line) ± SEM (dotted lines). Intensity traces were corrected for bleaching and aligned in time to the maximum intensity. (B) Same data displayed, but for cluster-like inclusions (N=4 CLIs).

A

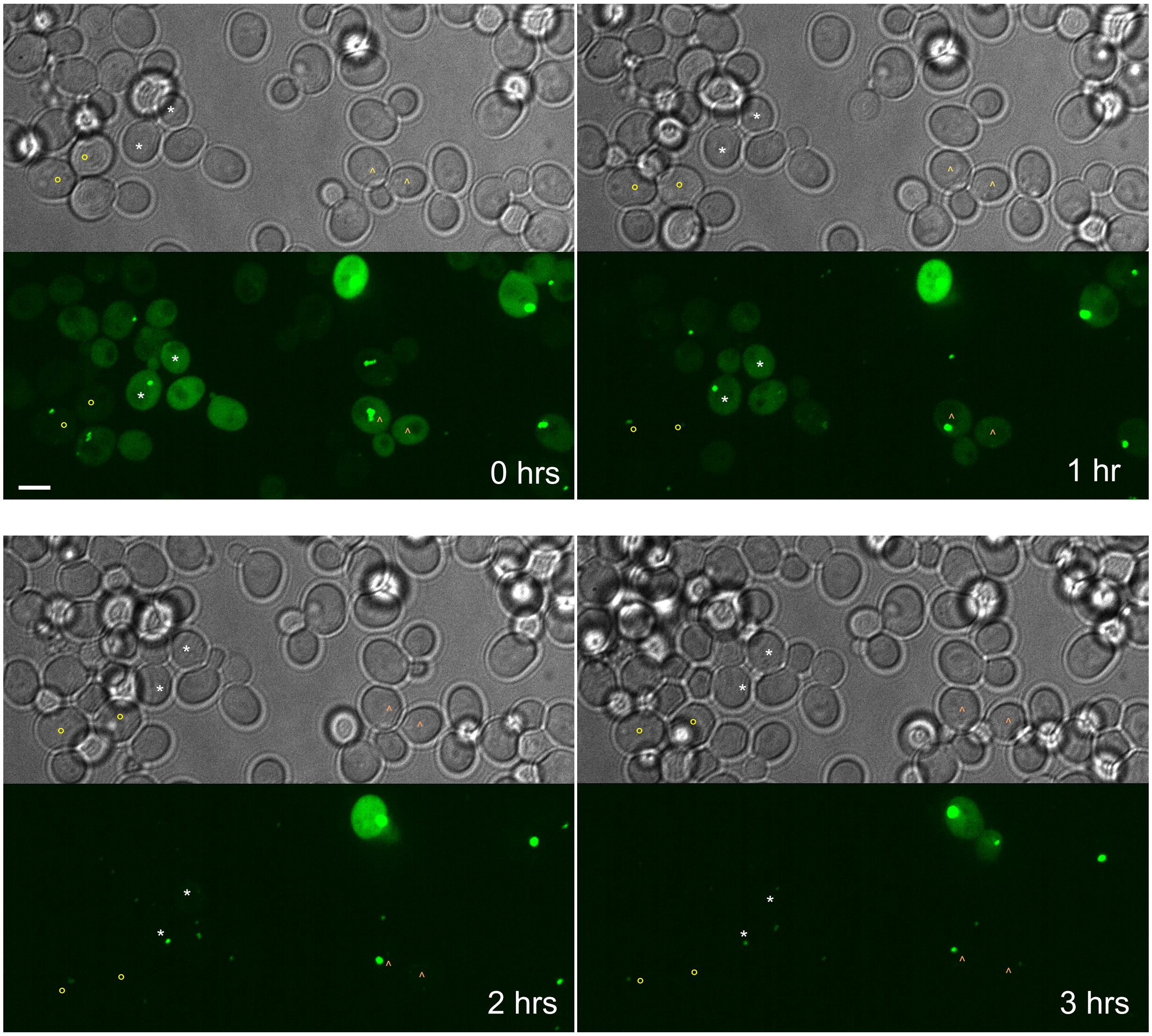


**Figure S5. Brightfield and fluorescent images of a field of mHtt-degron-GFP-expressing cells exposed to auxin**. A field of cells containing the 6 cells shown in Figure 6 is shown; maximum-intensity projections of confocal images (below) of with a brightfield image of the same field (above). The cells shown in Figure 6 are indicated with the symbols ° (yellow, left-hand column in Fig. 6), * (white, central column in Fig. 6), ^ (orange, right-hand column in Fig. 6), and. Scale bar, 2 μm.

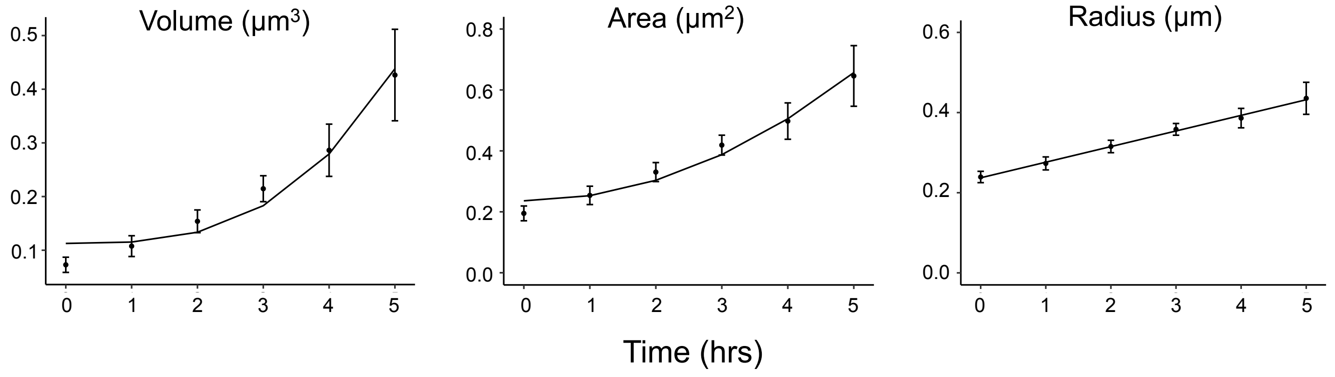


**Figure S6. IB growth rate for all IBs.** Average IB volume, area and radius were plotted over time for all IBs (error bars indicate SEM). Best-fit curves to functions of $t^3$ for volume ($R^2$ = 0.95), $t^2$ for area ($R^2$ = 0.97) and $t$ for radius ($R^2$ = 0.995) are shown.
