## Supplementary material for "Threshold concentration and random collision determine the growth of the phase-separated huntingtin inclusion from a stable core": Table S1

| **Quantity linear with time** | **Curve fit for radius vs** | | **Curve fit for area**  **vs** | | **Curve fit for volume**  **vs** | |
| --- | --- | --- | --- | --- | --- | --- |
|  | ***t^n^*** | **R^2^** | ***t^n^*** | **R^2^** | ***t^n^*** | **R^2^** |
| **Radius** | *t* | 0.99 | *t^2^* | 0.99 | *t^3^* | 0.97 |
| **Area** | *t^1/2^* | 0.84 | *t* | 0.95 | *t^3/2^* | 0.98 |
| **Volume** | *t^1/3^* | 0.69 | *t^2/3^* | 0.85 | *t* | 0.90 |

**Table S1. Curves fits for growth of IB with time.** For each model, linear growth of IB radius with time, linear growth of area with time, and linear growth of volume with time, curves were fitted to plots of radius, area and volume. For each model, the prediction of linear growth was extended to the other IB parameters: for example, if the growth of the radius with time is linear, it follows that the growth of area will grow as t^2^, and volume will grow as t^3^. Coefficients of determination (R^2^) for all curve fits of IB volume, area and radius with *t* are shown. Data for IBs in cells with cytoplasmic intensity >1100 was used (n=16).
