## Supplementary material for "Threshold concentration and random collision determine the growth of the phase-separated huntingtin inclusion from a stable core": Table S2

| **Cytoplasmic Intensity** | **Slope of**  **radius vs. time**  **(μm/hour ± SE)** | **95% confidence interval for slope** | **R^2^**  **For line fit** |
| --- | --- | --- | --- |
| 2695 | 0.071 ± 0.009 | 0.048 to 0.094 | 0.93 |
| 2654 | 0.075 ± 0.006 | 0.060 to 0.091 | 0.98 |
| 2181 | 0.052 ± 0.004 | 0.036 to 0.069 | 0.99 |
| 2130 | 0.061 ± 0.01 | 0.003 to 0.12 | 0.91 |
| 2087 | 0.060 ± 0.007 | 0.031 to 0.089 | 0.98 |
| 2010 | 0.047 ± 0.01 | 0.001 to 0.093 | 0.91 |
| 1981 | 0.038 ± 0.02 | -0.033 to 0.11 | 0.73 |
| 1857 | 0.039 ± 0.007 | 0.009 to 0.068 | 0.94 |
| 1782 | 0.062 ± 0.02 | 0.013 to 0.11 | 0.85 |
| 1686 | 0.054 ± 0.005 | 0.040 to 0.069 | 0.96 |
| 1616 | 0.034 ± 0.003 | 0.024 to 0.043 | 0.96 |
| 1445 | 0.041 ± 0.003 | 0.033 to 0.050 | 0.97 |
| 1392 | 0.044 ± 0.007 | 0.016 to 0.073 | 0.96 |
| 1329 | 0.029 ± 0.006 | 0.01 to 0.049 | 0.89 |
| 1272 | 0.021 ± 0.003 | 0.014 to 0.029 | 0.91 |
| 1234 | 0.049 ± 0.01 | 0.014 to 0.085 | 0.87 |
| 1091 | 0.025 ± 0.006 | 0.007 to 0.043 | 0.79 |
| 1058 | 0.020 ± 0.02 | -0.082 to 0.12 | 0.26 |
| 1029 | 0.027 ± 0.002 | 0.022 to 0.032 | 0.97 |
| 979 | 0.040 ± 0.008 | 0.013 to 0.067 | 0.88 |
| 859 | 0.021 ± 0.003 | 0.012 to 0.030 | 0.88 |
| 782 | 0.017 ± 0.008 | -0.008 to 0.042 | 0.61 |
| 743 | 0.022 ± 0.004 | 0.010 to 0.034 | 0.87 |
| 687 | 0.026 ± 0.01 | -0.011 to 0.063 | 0.62 |
| 504 | 0.0084 ± 0.005 | -0.0053 to 0.022 | 0.33 |

**Table S2. Slope of line fitted to a plot of radius vs time for individual IBs.** The cytoplasmic intensity of the cell in which the IB is found is given in the first column, the slope of the fitted line is given in the second column (best-fit ± SE). Goodness of fit is indicated by the 95% confidence interval and the coefficient of determination, R^2^.
