## Supplementary material for "Threshold concentration and random collision determine the growth of the phase-separated huntingtin inclusion from a stable core": Table S3

**Table S3** Plasmids

| **Plasmid** | **Description** | **Source** |
| --- | --- | --- |
| pEB4 | Htt(72Q)-GFP/CEN/LEU2, derived from p415GPD | 10 |
| pEB18 | Htt(72Q)-IAA^71-114^-GFP/CEN/LEU, derived from p415GPD | This study |
| OsTir1 | HO-Tir1 | 25 |
| GCN5-mCh | GCN5-mCherry- IAA^71-114^ | 25 |
