## Supplementary material for "Threshold concentration and random collision determine the growth of the phase-separated huntingtin inclusion from a stable core": Table S4

**Table S4** Strains

| **Strain** | ***Genotype*** | ***Source*** |
| --- | --- | --- |
| mHtt(72Q)-  GFP | *MATa his3Δ1 leu2Δ0 met15Δ0 ura3Δ0 pEB4 [pGDP-mHtt(72Q)-GFP::LEU2]* | Aktar et al., 2019 |
| HO-TIR1 | *MATa his3Δ1 leu2Δ0 met15Δ0 ura3Δ00 HO-TIR1* | This study |
| HO-TIR1 mHtt(72Q)-degron  GFP | *MATa his3Δ1 leu2Δ0 met15Δ0 ura3Δ00 HO-TIR1 pEB18 [pGDP-mHtt(72Q)-degron-GFP::LEU2]* | This study |
